# Biophysical Simulation Enables Multi-Scale Segmentation and Atlas Mapping for Top-Down Spatial Omics of the Nervous System

**DOI:** 10.1101/2024.08.02.605366

**Authors:** Lina Mohammed Ali, Aldrin Kay Yuen Yim, Emanuel Gerbi, Thien Nguyen, Nicholas Tu, Faith Ikede, Remi Sampaleanu, Diana Grigore, Jason Waligorski, Colin Kremitzki, Liya Yuan, Wendy Dong, Robi Mitra, Jeff Milbrandt, William Buchser

## Abstract

Spatial omics (SO) has produced high-definition mapping of subcellular molecules (like transcripts or proteins) within tissue samples. Mapping transcripts to anatomical regions requires segmentation, but even segmenting nuclei remains challenging for tissues like nerve cross sections, let alone for larger regions such as within the spinal cord. Neural networks could address this but need extensive human annotations—a bottleneck. We present SiDoLa-NS (**Si**mulate, **Do**n’t **La**bel – **N**ervous **S**ystem), an image-driven (top-down) approach to SO analysis in the nervous system. We utilize biophysical properties of tissue architectures to design synthetic images mimicking tissue samples. With these *in silico* samples, we train supervised instance segmentation convolutional neural networks (CNNs) for nucleus segmentation, achieving precision and F1-scores > 0.95. We take this a step further with generalizable CNNs that can identify macroscopic tissue structures in the mouse brain (mAP_50_ = 0.869), spinal cord (mAP_50_ = 0.96), and pig sciatic nerve (mAP_50_ = 0.995).

**Short Summary:** The SiDoLa-NS micro-, meso-, and macro-scale models are generalizable, supervised CNNs for neuronal segmentation in cell to tissue-level contexts. SiDoLa-NS is novel in its combination of three core ideas: it is top-down (image first), trained solely on synthetic images, and it is multi-scale. The tool is validated on brain, spinal cord, and sciatic nerve for advanced segmentation tasks.

## 1 Introduction

### 1.1 Images as Datasets

Technological advancements such as spatial omics have revolutionized our molecular mappings by enabling transcriptomic, proteomic, and metabolomic analyses within tissues. Yet, they fall short when it comes to leveraging image-derived features that capture the nuanced spatial organization of tissues. Questions about how molecular identities vary across tissue structures, for instance excitatory neuronal marker expression in the dentate gyrus versus the hypothalamus, remain hard to answer. Addressing such questions requires an integration of information across various scales: the cellular, regional, and tissue-wide scale. Simply put, what if you could automatically annotate structure? Thus, the need for tools that enable segmentation of tissue across these various scales and the contextualization of molecular patterns within the segmented regions.

### 1.2 Spatial Omics

Over the past decade, spatial omics (SO) technologies have revolutionized our ability to map transcriptomics, proteomics and metabolomics within histological sections, offering spatial localization insights at the tissue scale^1–9^. The immense throughput has provided transcript- and protein-wide data across diverse tissues including the spinal cord^10–13^ and mouse brain, an extraordinarily complex and dynamic region with specific architectural and physiological properties^8,14–19^. Addressing questions pertaining to spatial patterns of transcripts and proteins necessitates precise image segmentation and registration to biologically defined tissues^7,20,21^.

### 1.3 Bottom-Up Analyses

Most applications of SO are molecule-driven i.e. the spatial distributions of sets of transcripts are used to make distinctions between samples in the nervous system^22,23^. Such inferences do not automatically utilize brain atlas regions. The inclusion of spatial coordinates in omics datasets could unlock regional associations between RNA transcripts and proteins but are sometimes just an afterthought to the standard single cell pipeline. These ‘bottom-up’ approaches often employ clustering algorithms which learn locally and globally variable gene expression patterns^1,24–32^. GASTON models the transcriptomic space as a topographic map^33^, while Baysor treats cells as elongated transcriptional Gaussian distributions^34^. Bottom-up approaches like SpatialGlue and MIP-Seq^25,35^ integrate multi-omics data or cross-species expression^36^ but require multiple datasets. Unfortunately, using more genes to define regions reduces the number available for analyzing changes in expression patterns, leaving an insufficient pool for identifying unknown cell types^37,38^.

### 1.4 Top-Down Segmentation

Top-Down approaches determine regional information from image features or by alignment/registration. This methodology allows researchers to ask questions about molecular changes in a defined region and for cross validation of bottom-up transcriptomic analyses. Identifying pathologies and anatomical structures within tissue is a common use case. Original tools aimed to use interpretable algorithms based on local pixel values^39,40^ or registration^41,42^. Recent methods leverage deep learning^9,43–45^, transfer learning^46,47^, transformers^48–50^, and combinations to segment relevant structures and label cells or tissue types in immunofluorescence and histology datasets. These tools rely on either manual or autoencoder-based labeling, each presenting their own strengths and drawbacks. While autoencoders yield nuanced embeddings, they remove human oversight, degrading biological interpretability and noise sensitivity^51,52^. Human-labeled data enhances interpretability but increases the time to produce training sets, can introduce errors/biases, lacks widely available data^53^, and may hinder novel discoveries^54^. Despite this problem, there is a growing desire for tools that perform non-manual segmentation of biological data^55^. DeepSlice and FTUNet handle macro-scale atlas mapping and lesion segmentation, iStar and UniFMiR can enhance micro-scale data, and meso-scale analyses bridge tissue structures and the cells defining them^56,57^.

### 1.5 Multiscale Segmentation with SiDoLa-NS

What if you didn’t have to hand annotate regions to gain these insights or to train new models? In this manuscript, we present a tool for tissue segmentation across micro-, meso-, and macro-scales of cell and tissue organization. We use a biophysics engine to create image simulations to mimic tissue structurally at each level of interest: to identify nuclei and cell boundaries, to identify intermediate features (like clusters or bands of cells), and to identify regions with atlas mapping. Our *in-silico* image simulations offer 1) full control over feature distributions, 2) imperfection and noise representation (yielding tolerance), and 3) access to a virtually infinite and annotator-error-free training set. This is distinct from diffusion and GAN-based approaches, which take, as input, real microscopic images for synthetic data generation. Our datasets are used to train instance segmentation CNNs, resulting in SiDoLa-NS, which can then define cellular and regional identity based on image features alone and avoid biases from global gene expression and double dipping. We validate SiDoLa-NS with internal and publicly available brain, spinal cord, and sciatic nerve images and show the versatility of our tool across various platforms, including 10X Xenium, Visium SD/HD, H&E, and immunofluorescence sections. At the micro-scale, SiDoLa-NS performs cell and nuclei segmentation; at the meso-scale, it allows (for example) within- and between-fascicle analyses in the sciatic nerve; at the macro-scale, SiDoLa-NS defines brain regions based on the Allen Brain and Spinal Atlases^58–65^. SiDoLa-NS is a specialized top-down tool that automatically annotates nervous system tissue and gives researchers new ways to group the data across the scales and ask unbounded molecular questions: How do gene expression patterns of specific cell types vary across brain regions? What structural changes underlie molecular disruptions in disease? How do meso-scale features, such as clusters or bands of cells, relate to macro-scale organization in the nervous system?

## 2 Results

### 2.0 Overview of SiDoLa-NS

We present SiDoLa-NS (**Si**mulate, **Do**n’t **La**bel – Nervous System), a suite of tools designed for the detection and classification of image features within various contexts of the nervous system. SiDoLa-NS (pronounced See Doh La, think Do-Re-Mi) employs an image-based top-down approach, where labels are assigned to biologically defined regions of an image (**Fig. 1A**). This contrasts with frequently used bottom-up methods that cluster cells based on gene or protein expression, which can blur heterogeneous cell or tissue interactions. SiDoLa-NS models are trained on fully synthetic/simulated images, which are created with biophysical properties (physical properties of the tissue, such as cell density) *not* by generative AI (**Fig. 1B**). We use uniform distributions of parameters to build a variety of cells which fill regions, then convert these three-dimensional physical representations into images using an optics engine. We create images that reflect different staining modalities like immunohistochemistry (IHC) and immunofluorescence (IF). This has the advantage of providing truly massive datasets, where we can easily generate hundreds of thousands of images in a few hours. There is also no annotator bias, perfectly resolved ground truth segmentation masks and labels, and no memorization issues since the classifier never sees real images until inference. With the simulated images, we represent examples of various tissue and optical aberrations that may be present in a physical tissue section including 1) out-of-field cells, 2) broken tissue, 3) poor focus / contrast / resolution, and 4) distorted or warped tissue sections. We also over-represent examples that are difficult for existing segmentation algorithms, such as extremely high-cell density. Finally, ground truth labels and segmentation masks for the training images can be dynamically updated during simulation generation, enabling the entire schema to evolve—an impractical task if manually re-labeling 100K images. At no point during training are real biological samples used; both training and optimization utilize simulations, reserving the biological samples solely for downstream testing and analysis.

**Figure 1.**
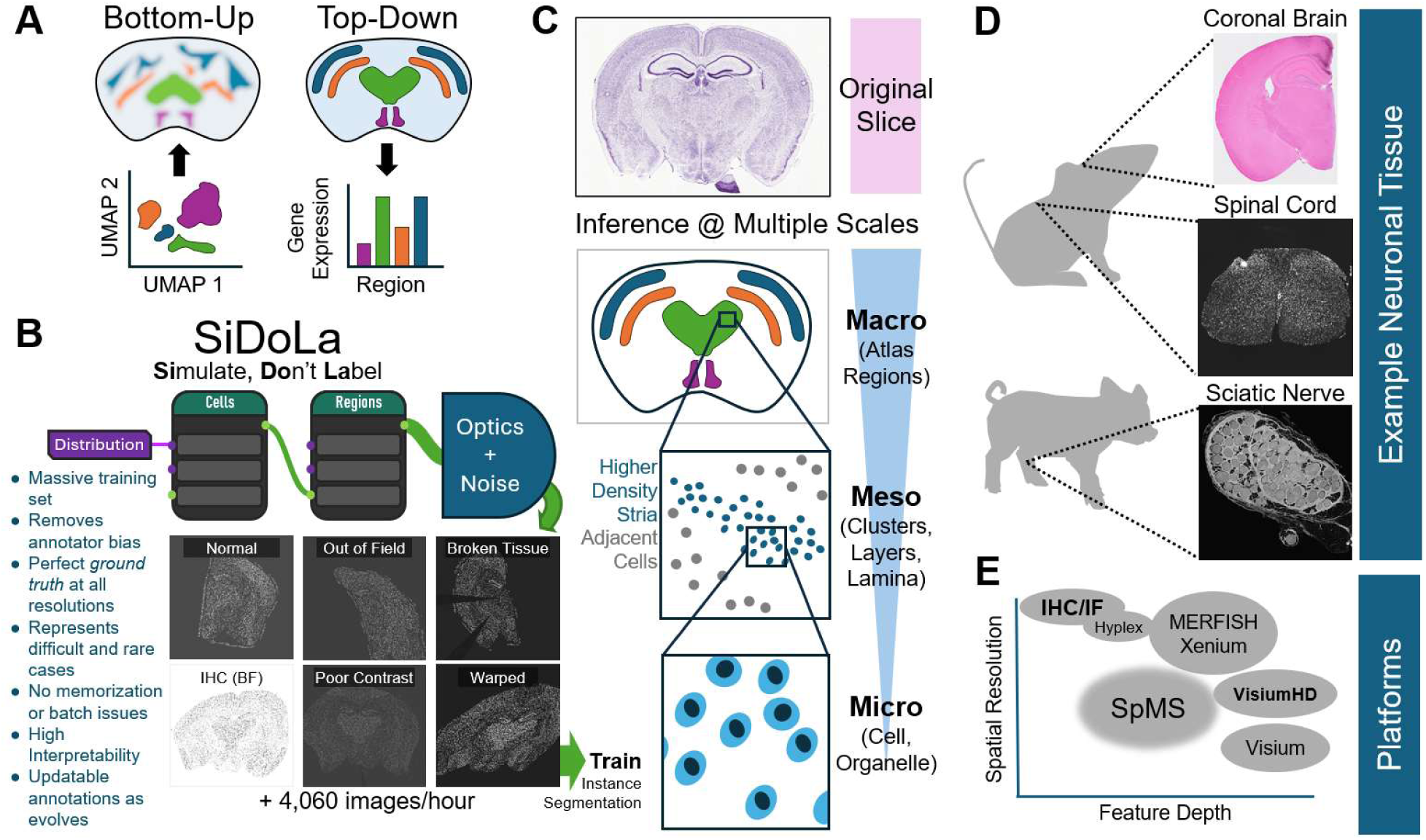
<u>Si</u>mulate <u>Do</u>n’t <u>La</u>bel: SiDoLa-NS for Top-Down Segmentation of Regions, Clusters, and Cells in Neuronal Tissue. **A)** Schematic illustrating the distinction between bottom-up and top-down methods. Top-Down approaches capture distinct image-based features and representations of a tissue. Bottom-Up approaches consider omics data (such as gene expression) to build molecular definitions of regions and cell types. **B)** Simulations create a massive synthetic dataset by stringing together biophysical nodes that build *Cells* and *Regions* of the image and then transform them from 3D geometry into images with an *optics* engine. Synthetic micrographs are shown below the nodes, giving examples of variation and features for a mouse coronal brain slice. **C)** Mouse coronal brain Nissl section from the Allen Brain Atlas, shown as the “original slice”, with cartoons below and the concept of multiple spatial scales where segmentation identifies individual instances of a variety of features. Specifically, this approach identifies atlas-based regions at a <u>macro</u>scopic scale, as well as cell density and other <u>meso-</u>scale patterning in the tissue and is also able to segment cells to incorporate subcellular <u>micro-</u>scale information, like cell type and structure. **D)** SiDoLa-NS was tested on three nervous system tissue types including mouse brain and spinal cord, as well as porcine sciatic nerve (a human-scale nerve cross section). **E)** These tools are designed to work with the full spectrum of image-based datasets from classical immunohistochemistry to spatial transcriptomics platforms.

While each model is employed at a specific scale, SiDoLa-NS streamlines the integration of data across multiple scales. This is useful for tissue such as the brain, where it is essential to understand the spatial extents of regions from a brain atlas, to find clusters, lamina, and striations of cells, and to carefully detect nuclei (**Fig. 1C**). In this manuscript, we examine the results of SiDoLa-NS against three publicly available datasets from mouse and porcine neuronal tissues, including the brain, spinal cord, and sciatic nerve (**Fig. 1D**). SiDoLa-NS is designed to work with a wide range of platforms, from classical IHC and IF tissue sections, to the full range of spatial omics technologies (**Fig. 1E**). Lower resolution spatial omics, like the original 10X Visium platform, still operate normally since the accompanying tissue section imaging is done at relatively high resolution, even if the transcript data is obtained at a lower one.

### 2.1 SiDoLa-NS is Trained on Biophysical Simulations of Nuclei in Brain Slices to Segment a Histological Section of Mouse Brain

We first asked whether we could segment cells, layers, and atlas regions within a mouse brain slice. We constructed biophysical representations of mouse brain sections at multiple scales. The first fully synthetic image set (with 4,353 pairs and 1.96×10^6^ objects) was at the cellular level and mimicked the DAPI staining phenomena observed in the nuclei of cells in the brain (**Fig. 2A 1**). These synthetic images represented neuronal, oligodendrocyte, and other nuclei of different size, shape, and intensity.

**Figure 2.**
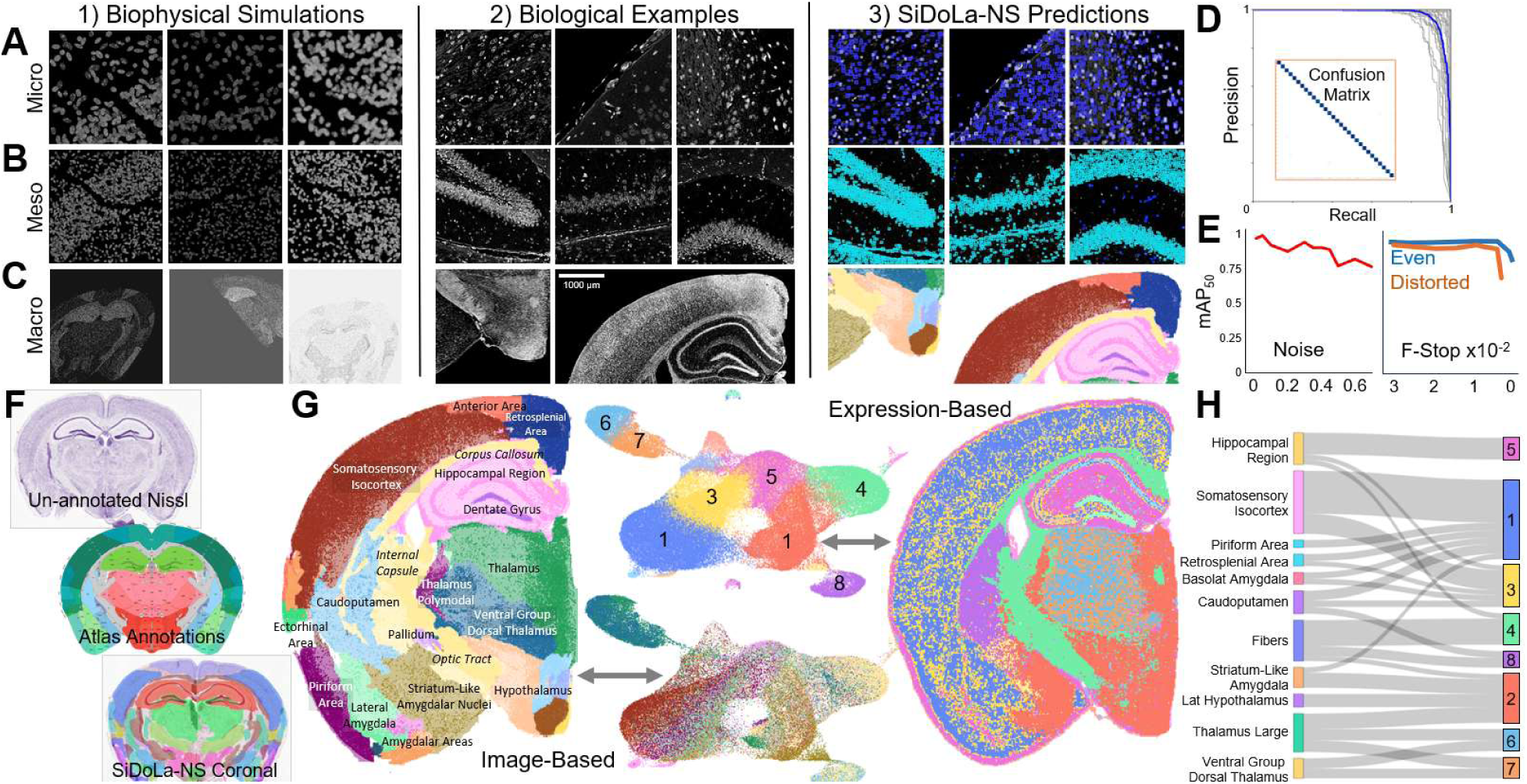
Multi Scale Segmentation and Classification of a Mouse Coronal Brain Section. **A)** At the micro-scale, SiDoLa-NS trained exclusively on simulated data (1) featuring multiple cell types. Micrographs of real brain images (2) were used to run SiDoLa-NS-Micro-CNS at this scale and identified nuclei (light and dark blue bounding boxes) from background (3). **B)** At the meso-scale, simulated images (1) featuring bands of higher density cells. Micrographs from brain slice, notice density patterns (2). SiDoLa-NS classes overlaid, which differentiated high density striated regions (cyan) and adjacent cells (blue) (3). **C)** At the macro-scale, SiDoLa-NS was trained on simulated images (1) featuring full or hemi coronal sections of brain tissue. SiDoLa-NS-Macro-mCB segmented real images of an H&E coronal brain slice (2), highlighting biologically defined regions (3). **D)** Precision-recall curve and confusion matrix for the macro atlas predictions. **E)** Line graph that depicts mean average precision (mAP_50_) on the y-axis vs. different amounts of image manipulation on the x-axis. The left panel increases white noise proportion from 0 to 0.7. The right panel increases blur by lowering the aperture’s f-Number in both even and distorted (blur was differential across the image) cases until ∼0.01, where mAP_50_ dropped. **F)** Unannotated, Nissl-stained mouse P56 coronal slice, alongside Allen Brain Atlas annotations and SiDoLa-NS annotations (the Nissl brain is 9730 µm in width). **G)** Comparison of Top-Down and Bottom-Up approaches side-by-side. The left-brain image and lower UMAP are SiDoLa-NS top-down, while the right brain and upper UMAP are bottom-up. The UMAP was constructed from transcriptomic data, and its colors are by Seurat clusters in the upper UMAP, and by SiDoLa-NS labeled brain regions in the lower. **H)** Sankey diagram mapping SiDoLa-NS-Macro-mCB brain regions to the Seurat clusters in Figure 2G.

Renderings intentionally spanned a wide set of poorly- and well-focused images. We then trained an *instance segmentation* CNN to find these nuclei, resulting in SiDoLa-NS-Micro-CNS. For prediction, we used a publicly available 6mm x 6mm H&E (hematoxylin and eosin) coronal cross section of mouse brain which was run on 10X Genomics’ Visium HD platform. Zooming in on small tiles, the full brain image showed varied morphology from a single channel of the H&E staining (**Fig. 2A 2**). The results of inferring on the biological examples with SiDoLa-NS-Micro-CNS trained on simulated images are shown, where the brighter and dimmer nuclei are both detected, but given different class labels (**Fig. 2A 3**). Increasing the perspective to see larger numbers of cells, but still not the whole brain, we refer to as the “meso-scale”. Here we observed two cellular populations with distinct densities, which we refer to as ‘striated’ or ‘laminar’ regions. These striated regions are bands of high nuclear density, whereas adjacent laminar regions were observed with more diffuse nuclei (**Fig. 2B 1**). Striated regions include the granular cell zones (for example of the Dentate Gyrus or Hippocampal CA regions), which are composed primarily of high-density neuronal cell bodies (**Fig. 2B 2**) and were detected (**Fig. 2B 3**) in the Visium HD brain slice. Model performance was measured with the synthetic validation dataset. SiDoLa-NS-Micro-CNS achieved high IoU (Intersection over Union) scores for detection and an mAP_50_ (mean average precision for IoU greater than 0.5) of 0.715 (**Fig. S1**).

We also asked whether SiDoLa-NS-Micro-CNS could work beyond the nervous system, therefore testing its generalizability. We chose a publicly available DAPI stained image of lung adenocarcinoma (10X Genomics Visium HD), which is challenging for other segmentation algorithms due to regions of densely packed cells. We observed excellent segmentation and detection of nuclei, even in high-density regions of this tissue (no parameters changed from brain inference), demonstrating that the network is generalizable and unlikely to succumb to issues with batch effects (**Fig. S2**).

### 2.2 SiDoLa-NS is Trained on Biophysical Simulations of Brain Slices and used to Map Brain Regions in a Histological Section of Mouse Brain

At the macro-scale (where the entire section of brain is visible), we asked whether we could assign named regional identities to the detected nuclei. To accomplish this, we created a dataset of fully synthetic coronal slices of the mouse brain (single hemispheres and full). These macro-scale simulations were built from region labels based on the Allen Brain Atlas reference for the mouse brain (**Fig. 2C 1**). We used 144,438 training images and 1,092 validation images (all synthetic) to train the CNNs, which can be combined in an ensemble. Using the same coronal brain slice (**Fig. 2C 2**), SiDoLa-NS inferred brain regions which clearly delineate distinct brain areas (**Fig. 2C 3**). The primary CNN (SiDoLa-NS-Macro-mCB) achieved an excellent precision-recall curve and the confusion matrix on the validation set was a near-perfect diagonal, indicating close to zero class confusion (**Fig. 2D**). We measured a top mAP_50_ of 0.977 on the validation set (**Fig. S3**).

Another feature of the SiDoLa-NS suite is their robust performance even with noisy or blurry samples. To demonstrate SiDoLa-NS-Macro-mCB’s ability to handle these imperfections, we inferred on images where noise was introduced in a calibrated fashion (see Methods) and found that SiDoLa-NS maintained an mAP_50_ above 0.783, even when most of the image was noise (up to 70% noise mix, **Fig. 2E**). Next, we simulated images where focus was reduced across the entire image (even) or in certain parts of the image (distorted). SiDoLa-NS-Macro-mCB maintained an mAP_50_ above 0.803 and 0.722 across the even and distorted images respectively (where the aperture f-number was reduced to as low as 0.0013).

While it was clear that the model was robust to noise and blur, and comparable to other algorithms with similar function, we wanted to validate that these networks maintained high performance even when applied to real biological images. We specifically sought to test versatility with *staining modality*, and we inferred on unseen Nissl-stained brain samples with corresponding true labels, provided by the Allen Brain Atlas (**Fig. 2F**). Here again, even though the Nissl image was of the whole brain and had a bright background (the colors were not inverted prior to inference), SiDoLa-NS was still able to find the brain regions and segment them.

We were interested in seeing the differences between a top-down vs a bottom-up approach to cellular labeling. Specifically, we tested how SiDoLa-NS would look relative to Visium HD’s Seurat pipeline for clustering and spatial overlay. We inferred SiDoLa-NS-Macro-mCB on the whole mouse brain slice which automatically links these region labels to the cell-level segmentations and transcripts. Here, both individual cells and polygonal areas are assigned brain regions (**Fig. 2G**). We compared the atlas predictions (Top-Down) to the transcriptomic data (Bottom-Up) directly by overlaying the predicted region labels over the Seurat-generated UMAPs, or by overlaying the Seurat clusters over the brain image. As expected, the two methods are complementary, with the top-down excelling at finding boundaries and filling the whole area, while the bottom-up at finding different cell types within regions. For example, cells which SiDoLa-NS assigned to the Striatum-Like Amygdala, Lateral Hypothalamus, and Thalamus were difficult to differentiate by the molecular cluster. Using both together can help disambiguate numbered clusters and match up to named atlas regions (**Fig. 2H**).

We sought to compare our image-based regional definitions against expression-based regional definitions using Kaur et al’s^66^ consensus tissue domain detection on a publicly available mouse coronal (standard) Visium dataset. This yielded four sets of labels to compare with SiDoLa-NS-Macro-mCB atlas labels. All the methods used the expression data and variations of unsupervised clustering to define numbered discontinuous spatial regions. This comparison is biased towards the expression techniques, since we compared all the labels back to the molecular definitions of the regions from the consensus tissue domain manuscript. The comparisons (**Fig. S4**) are against MILWRM (multiplex image labeling with regional morphology), SpaGCN (spatially variable genes by graph CNN), and GraphST (Graph Spatial Trans)^24,27,66^. We found that MILWRM performed the best, with SpaGCN, and SiDoLa-NS close behind (accuracy 69.4%, 63.9%, and 60.1%) with GraphST having the most differential patterns as defined by MILWRM from expression (accuracy 51.7%). Additionally, we found that the molecular-defined labels incorrectly include hindbrain and olfactory bulb in a mid-coronal slice. This suggests that expression-based methods suffer from a lack of anatomical resolution and are more prone to transcript region misidentification.

### 2.3 SiDoLa-NS for Spinal Cord Nuclei Detection and Atlas Mapping

We next asked whether a less-commonly studied neuronal tissue, the spinal cord, could be analyzed with this system. SiDoLa-NS was evaluated on publicly available immunofluorescence (IF) sections of mouse spinal cord stained with DAPI for nuclear localization, NeuroTrace (NT) for neuronal soma, and ChAT for motor neurons^64^. As a side note, this mouse spinal cord slice was not a spatial omics dataset, and therefore lacked transcriptional data. At the micro-scale, we used SiDoLa-NS-Micro-CNS (**Fig. 3A 1**), even though the spinal cord IF section was lower resolution and slightly blurrier (**Fig. 3A 2**) than in the brain.

**Figure 3.**
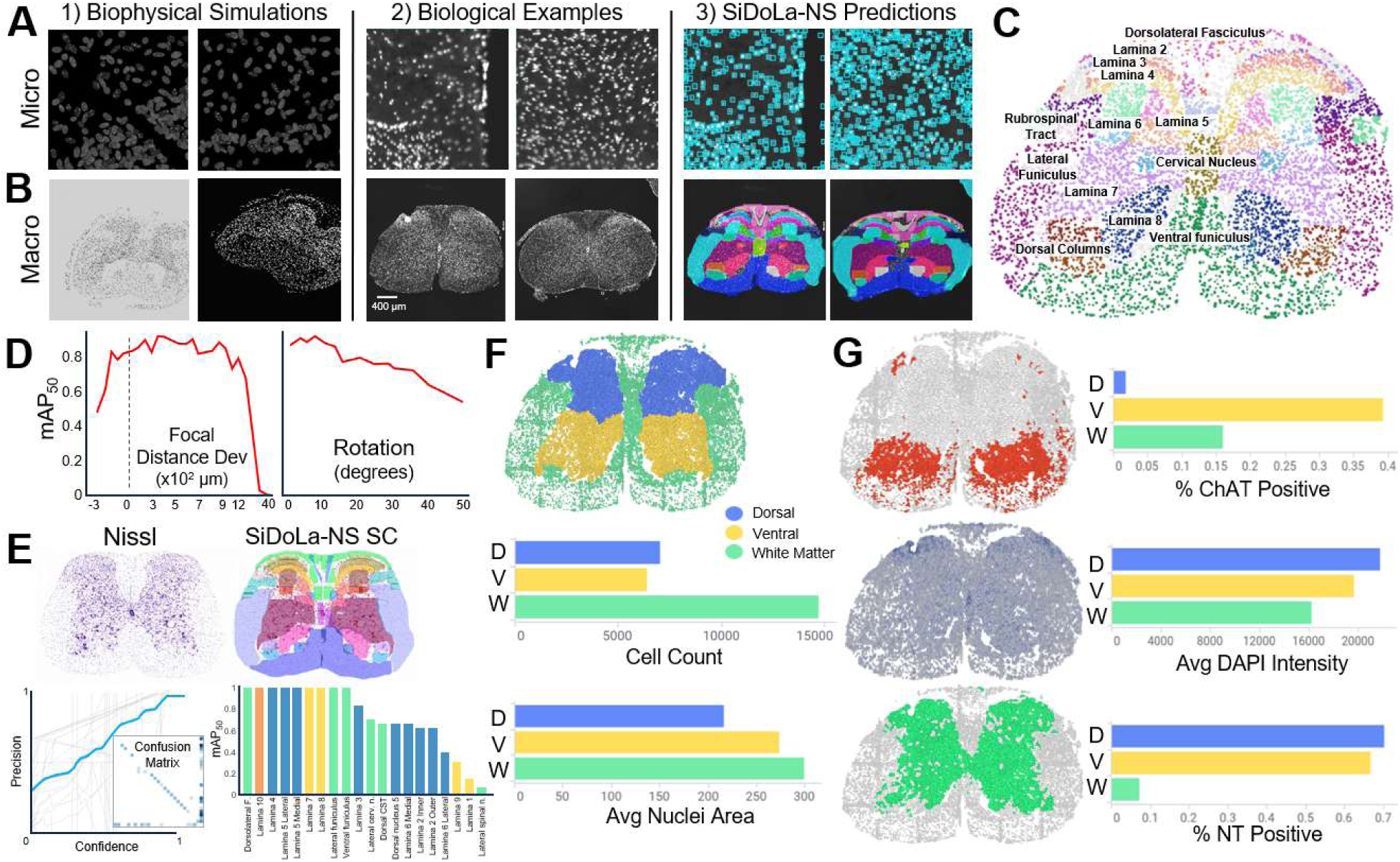
Segmenting Nuclei and Mapping Atlas Regions in a Mouse Spinal Cord Section. **A)** At the micro-scale, SiDoLa-NS trains on simulated data (1). Micrographs of tiles from cross sectioned spinal cords from dataset^64^ (2). SiDoLa-NS-Micro-CNS predictions (blue squares) of nuclei in the spinal cord (3). **B)** At the macro-scale, SiDoLa-NS trains on simulated images of spinal cord sections (1). Micrographs of mouse spinal cord cross sections. SiDoLa-NS-Macro-mSC prediction on different spinal cord regions (different colors) (3). **C)** Cellular and regional results on the whole spinal cord section. Marker size is governed by the measured cell size, and coloration is by regions, some of which are labeled. **D**) Line graph indicating performance of SiDoLa-NS-Macro-mSC at different focal distances and tilts. Y-axis shows mean average precision at an IoU of 50%. The left x-axis measures the delta focal distance, where the dashed line is centered at 0, the normal camera distance, and the spinal cord takes up the majority of the frame. As the focal distance increases, the camera is backed away making the image smaller in the frame. The right x-axis displays degrees of tilt along the y-axis, rotating the camera around to the right side, skewing the sample. **E)** Micrograph of a Nissl section from the Allen Brain Mouse Spinal Atlas (section #875) on the left, and the same image with SiDoLa-NS-Macro-mSC predictions overlayed (different colors are different regions). Below left, line graph of a Precision/Confidence curve, all classes at 0.96 precision with a confidence of 0.993. Inset, confusion matrix, and bottom left, a bar chart showing the mean average precision. All comparing SiDoLa-NS-Macro-mSC to Allen Brain Atlas defined regions. **F)** Cellular representation of the same spinal cord section from C) but colored to define 3 main regions within the spinal cord (blue dorsal horn, yellow ventral horn, and green is white matter). Below are bar charts depicting the cell count and average nuclei area per region. **G)** The cellular spinal cord representation, but now the cells are colored to depict, on the top: ChAT+ cells in red, with the bar chart measuring the % of cells positive in each region, middle: the intensity of DAPI staining (higher intensity is a darker shading), and bottom: NeuroTrace (NT)+ cells in green.

Still, SiDoLa-NS-Micro-CNS with identical settings worked to segment the nuclei (**Fig. 3A 3**). Utilizing the model, we found sufficient generalizability to the mouse spinal cord nuclei images. This demonstrated the robustness and versatility of SiDoLa-NS across tissues and, more challenging, across platforms and image modalities.

At the macro-scale, we simulated whole and hemi spinal cord histological slices and varied parameters including noise, warp, and tissue degradation (**Fig. 3B 1**). Spinal cord atlas mapping, like in the brain, previously required manual image registration; including these distortions in the SiDoLa-NS-Macro-mSC training set allowed for streamlined application on this tissue section. Using this macro-scale simulated image set (12,609 training, 30 validation), we trained SiDoLa-NS-Macro-mSC for whole slice spinal cord atlas mapping in mouse. We then inferred this model on biological IF whole tissue sections (**Fig. 3B 2**), highlighting the various spinal lamina alongside other prominent regions in the white and grey matter (**Fig. 3C 3**). SiDoLa-NS-Macro-mSC achieved a top precision of 0.96 at a confidence of 0.92 on the validation set (**Fig. S5**). We next examined its robustness to variations in focal distance and tissue rotation. SiDoLa-NS-Macro-mSC was resistant to focal distance variations (zooming in to -300 µm and out to 4000 µm from a normal distance of 675 µm between the sample and the camera); the CNN maintained an mAP_50_ above 0.70 within a range of -200 µm to +1000 µm from the standard distance used in the training set images (**Fig. 3D**). Zooming in too tightly quickly degraded performance presumably because it lost the spatial context provided by the boundaries of the spinal cord. Similarly, we tested the model’s robustness to degree of rotation, and found that in our test case, SiDoLa-NS-Macro-mSC maintained an mAP_50_ above 0.60 up to 40 degrees of rotation along the y-axis (tilting one side of the plane of the image ‘towards the camera’).

We inferred SiDoLa-NS-Macro-mSC on an unseen Nissl-stained image to examine the model’s abilities under varying tissue and staining conditions. SiDoLa-NS-Macro-mSC achieved a top precision mAP_50_ of 0.96 at a confidence of 0.929 (**Fig. 3E**). The mAP_50_ was class-variable, where larger and more well-bounded regions like the dorsolateral fasciculus achieved precisions of 0.995 and the thinner layers, like Lamina 1 and 2, had lower precisions (0.145 and 0.622 respectively). This is likely due to slight shifting of where the Allen brain polygons occurred compared to where the predicted segmentation masks were placed.

To validate the regional assignments on the IF mouse spinal cord sections, we compared the staining intensities in the reported atlas regions. When inferring, SiDoLa-NS multiscale results provide quantitative measurements for each predicted object at the micro-, meso-, and macro-scales, including area, channel intensity, and texture (measured as standard deviation of pixel intensities within the object). We combined the SiDoLa-NS-Macro-mSC predicted regions to three generic groupings: white matter, dorsal, or ventral gray matter. As expected, regions labeled as white matter were located on the outer perimeter of the spinal cord, enclosing dorsal and ventral regions. Dorsal gray matter occupied the upper half of the spinal cord, and ventral regions fell in the lower half (**Fig. 3F**). We next quantified the variable feature distributions across these groupings. For example, the white matter regions had slightly larger nuclei area. After applying an intensity threshold, we examined ChAT-positive cell distribution (**Fig. 3G**). We expected the ventral horn to contain the greatest fraction of ChAT-positive cells and indeed, the ventral horn was 38% ChAT+ compared to only 2% in the dorsal horn. As expected, we observed less than 10% NeuroTrace-positive cells in the white matter (compared with 67 and 71% in the ventral and dorsal horns, respectively) and a uniform nuclear intensity.

### 2.4 Axonal and Nuclear Segmentation within Sciatic Nerve Fascicles

Few resources exist for the segmentation of peripheral nerves. We sought to detect axons and Schwann cell nuclei as well as the fascicles that contain them within a cross section of a human-sized sciatic nerve. Here, we evaluated SiDoLa-NS’s ability to detect axons, nuclei and meso-scale features in the nerve. In both H&E and DAPI staining, it is possible to discern a distinction between axons, which are larger, lower intensity structures, and the nuclei of Schwann cells, which are smaller and brighter. Thus, we created simulations (7,198 training and 612 validation images, 2.78×10^5^ objects in the training set) at the cellular (micro-) scale with simple geometry (**Fig. 4A 1**). We trained SiDoLa-NS-Micro-PNS to explicitly segment axons and nuclei and distinguish them as unique classes. The model achieved a perfect precision at a confidence of 0.894, and an F1 score of 0.9 at a confidence of 0.4 on simulated validation images (**Fig. S6**). We next used a publicly available cross section of a porcine sciatic nerve, which is composed of bundles of axons and glial nuclei (**Fig. 4A 2**). When inferring SiDoLa-NS-Micro-PNS on the nerve, it detects the axons and nuclei within the tissue section (**Fig. 4A 3**).

**Figure 4.**
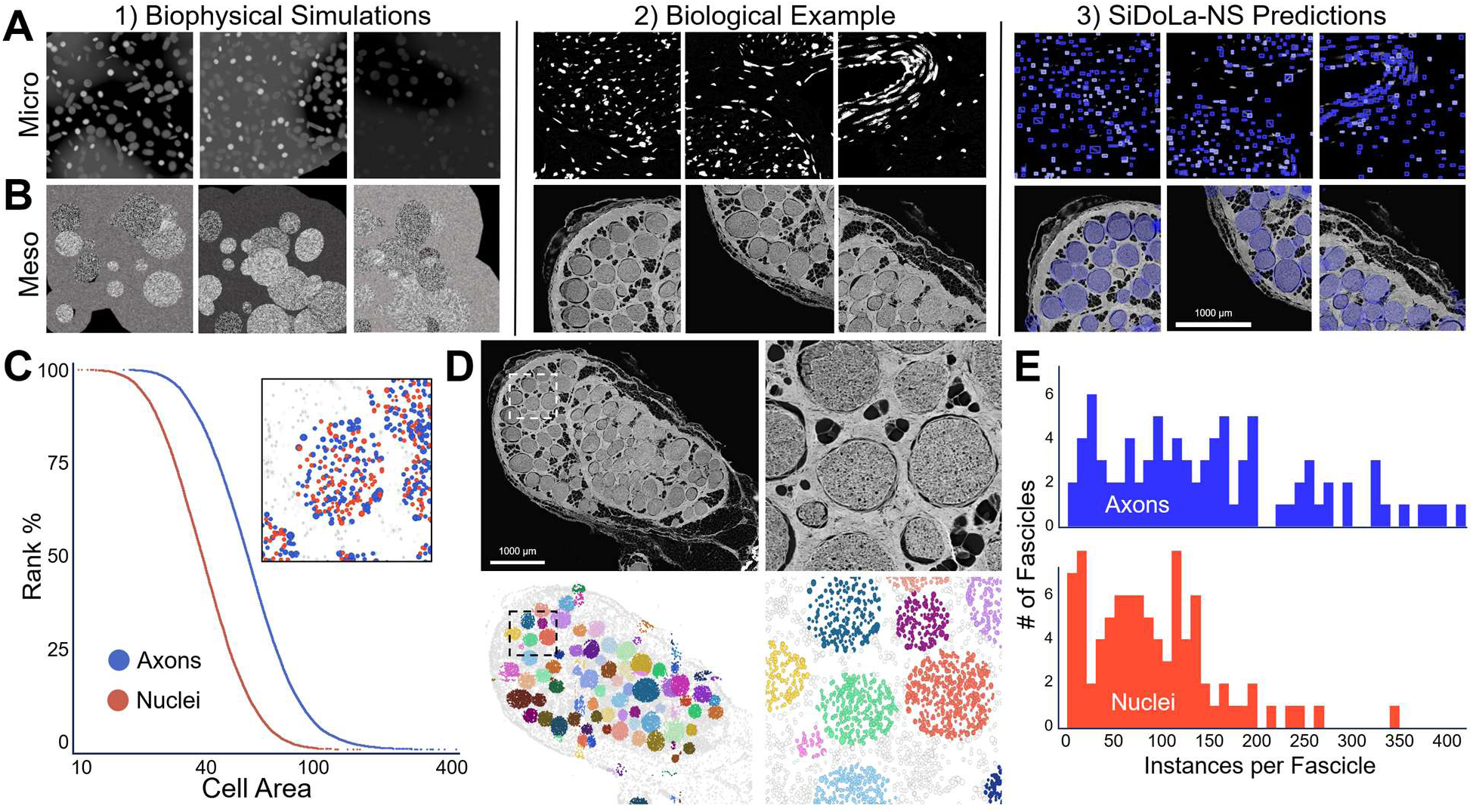
Porcine Sciatic Nerve with Automated Fascicle Identification and Axon/Nuclei Discrimination. **A)** Micrographs of a simulated cross section (1) of sciatic nerve with high intensity nuclei and dim axons. (2) Example tiles of a cross section of a porcine sciatic nerve and (3) SiDoLa-NS-Micro-PNS prediction on axons (dark blue) and nuclei (light blue) on the same sections. **B)** Micrographs of simulated images (1) of textured bundles (fascicles) and perineurium using varying background contrast. (2) Micrographs of a cross section of a porcine sciatic nerve. (3) SiDoLa-NS-Meso-pSN prediction of fascicles on porcine sciatic nerve section (blue circles). **C)** Ranked histogram showing distribution of axon (blue line) and nuclei (red line) relative pixel areas within all regions of the porcine sciatic nerve section shown. Inset is a small region of the nerve showing non-fascicle-contained cells (gray), axons (blue), and nuclei (red). **D)** Micrograph showing overview of entire sciatic nerve, with SiDoLa-NS-Meso-pSN fascicle detection below. The dashed box indicates the area shown in higher magnification on the right. Circular markers are axons or nuclei, and their color indicates different fascicle instances. **E)** Histograms depicting the cell contents within fascicles. The blue histogram represents axons, and the red histogram represents nuclei. Fascicles with >= 10 are shown except for one outlier with 740 axons not shown.

While mouse, human, and pig have little difference at the cellular scale inside a sciatic nerve, the peripheral nerves in the larger mammals are composed of many fascicles (bundles) packed with axons and Schwann cells and are surrounded by perineural and epineural tissue. Most neurodegenerative diseases are specific to one class of neurons, for example sensory neurons in CMT2A^67^ or motor neurons in ALS^23^, and the fascicles are impacted differentially due to their composition. We asked whether we could automate fascicle segmentation and assign the cells inside to specific fascicles. We created meso-scale simulations for images of fascicles as ellipsoid objects with varying size, degree of overlap, and fascicle/background intensity contrast (**Fig. 4B 1**) with 6,588 training and 304 validation images (89,074 training objects). This was then trained with instance segmentation, and the result was termed SiDoLa-NS-Meso-pSN. On simulated validation images, this model performed even better, with F1 reaching 1.0 for confidence values between 0.1 and 0.9 (**Fig. S7**). On the section of porcine sciatic nerve (**Fig. 4B 2**), single fascicle predictions were identified as ellipsoids in the tissue (**Fig. 4B 3**).

Next, we inferred on the porcine sciatic nerve section on the micro- and meso-scales, where nuclei, axons, and fascicles are all fused in one resulting dataset. At the cellular level, we confirmed the expected variation in the cell area between axons and Schwann cells, where axons are larger on average (**Fig. 4C**). At the meso-scale, SiDoLa-NS-Meso-pSN detected fascicles throughout the tissue slice, and unique fascicles were identified as numbered ‘instances’, where each cell within also had a feature indicating to which fascicle instance it belonged (**Fig. 4D**). We next asked what the variation in the number of axons or nuclei was in each fascicle and found that most porcine sciatic nerve fascicles had close to 100 axons, and about 70 nuclei, consistent with hand counting (**Fig. 4E**). Additionally, we noticed that fascicles on the lower left of the section had more nuclei, while the ones on the top right tended to have a lower percent of nuclei (data not shown). Just from an H&E nerve section, SiDoLa-NS-Meso-pSN allowed for basic analyses of the variation between and within fascicles.

### 2.5 Interrogating RNA Expression patterns and Cellular Morphology using Top-Down Analysis

Thus far, gene expression patterns have been used only to validate SiDoLa-NS’s predictions at the macro- (reference) scales. The top-down analysis after SiDoLa-NS inference allows for downstream investigations via measuring features across the SiDoLa-NS predicted regions. These features can be used, for example to define marker genes for distinct tissue regions. It can also be used to better compare regional expression changes between case and control sections.

In the mouse brain, the 10X Visium HD expression data was linked to each cell, and we interrogated RNA expression patterns across cells and regions predicted by SiDoLa-NS-Micro-CNS and SiDoLa-NS-Macro-mCB, respectively. We identified a list of region-specific genes, ones that had the highest ratio (region_x_’s average transcript count divided by the average transcript count of all regions excluding *x*) and checked their expression across the atlas regions **(Fig. 5A**). Most of these genes were also quite specific (strong signal along the diagonal), but a few had some off-diagonal hotspots between related regions. For example, *Ltih3*, *Calb2,* and *Agt* were all differentially expressed in the hypothalamus and hypothalamic-adjacent regions. We also compiled a list of the top 4 highest ratio genes (**Fig. 5B**), highlighting known markers such as *Tshz2* in the retrosplenial area and *Neurod6* in the hippocampus in addition to less commonly used marker genes like *Prkcd* in the ventral group of the dorsal thalamus. A larger, hierarchically clustered heatmap shows additional genes of interest per regions (**Fig. S8**). As expected, these genes maintain distinct regional localization in the coronal mouse brain (**Fig. 5C**). Additional gene localization patterns for *Tac1*, *Penk*, *Mobp*, *Acta2*, and *Prox1* were also visualized^68^ (**Fig. S9**).

**Figure 5.**
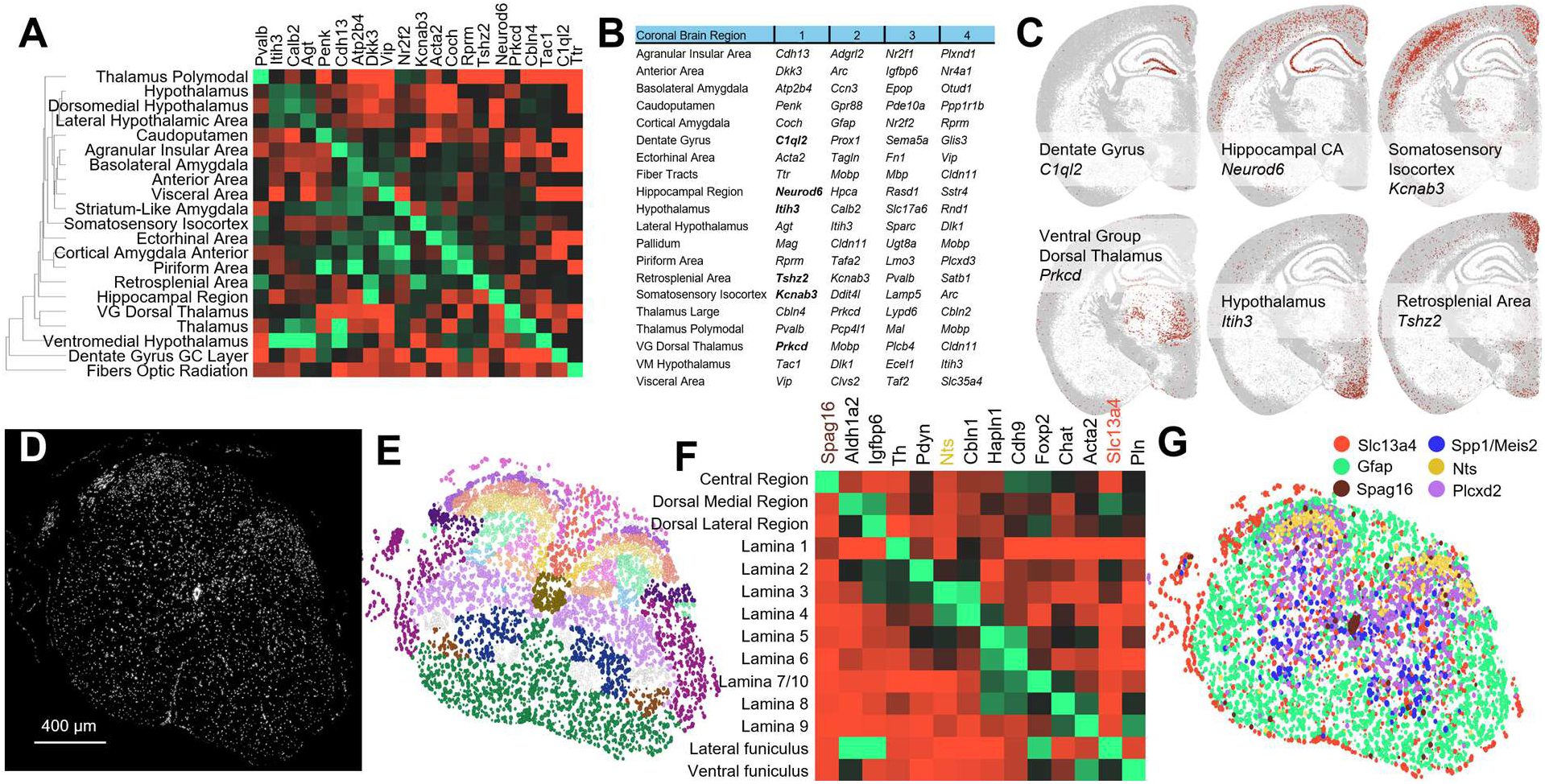
Top-Down Anatomical Regions Allow Unbiased Transcriptomic Questions in Mouse Brain and Spinal Cords. **A)** Heatmap showing the expression of the ‘top ratio’ genes for 21 regions from the mouse cortical Visium HD brain slice (green is high average counts, red is low). **B)** A list of the top four genes per brain region, where each gene was ranked by ratio of average count per region compared to the average counts in all other regions. Bolded genes are detailed in the next panel. **C)** Diagram of individual cells segmented with SiDoLa-NS-Micro-CNS, colored by total counts per cell for the gene specified (gray = 0, black = mean, red = max. **D)** DAPI-stained transverse lumbar spinal cord section. There is some tissue delamination around the edges and a clockwise rotation of the slice (Scale bar 400 microns). **E)** Cellular and regional results on the transverse section in D. Markers correspond to cells, marker size is governed by the measured cell size, and coloration is by regions. Low confidence (< 0.3) results discarded. **F)** Heatmap showing the expression of the ‘top ratio’ genes for 14 regions from the mouse spinal cord slice (green is high average counts, red is low). Some regions were grouped for clarity: Central Region contains Lateral Cervical Nucleus and Central Canal; Dorsal Medial Region contains Dorsal corticospinal tract, Dorsal Nucleus, and Dorsolateral fasciculus; Dorsal Lateral Region contains Rubrospinal tract and Lateral spinal nucleus; Lamina subgroups were merged, including Lamina 10 with the Lamina 7 group. Highlighted genes appear again in subpanel G. **G)** Some selected top hits from Kruskal-Wallis analysis were used to color the slice. Colors represent which of the 6 selected genes each cell expressed in the highest proportion. Spp1/Meis2 marked cells co-express these two genes.

The same process was performed on 10X Xenium expression data for mouse spinal cord (**Fig. 5D**). SiDoLa-NS-Macro-mSC identified the same regions noted in Fig. 3C (**Fig. 5E**), though notably this slice came from a distinct lumbar section and animal. A heatmap was generated following the same specifications as in Fig. 5A, though some subregions were merged to aid in visualization (**Fig. 5F**). Like Fig. 5A, many regions showed specific genes at higher expression: *ChAT*, traditionally associated with motor neurons was most highly expressed in Laminas 8 and 9, as expected^69^, and *Spag16*, which codes for a building block of cilia, was located around the central canal, where the cerebrospinal fluid is propelled largely by these microscopic hair-like motors^70^. Although the spinal cord is largely neural tissue, we considered that epithelial and muscular tissues may also impact analyses of this nature. For instance, *Acta2*, a gene associated with smooth muscles and found in vasculature throughout the CNS^71^, defined lamina 9 in this analysis, an incidental finding. Adjacent regions often showed overlapping expression, like Lamina 5 and 6. Additionally, the lateral funiculus, which contained delaminated tissue from the extreme left and right sides of the slice, had multiple highly expressed genes. Kruskal-Wallis was performed on the spinal atlas regions and Xenium expression data to rank genes by differential expression. (**Fig. 5G**). We selected a handful of the spatially segregated genes from this analysis (some of which also appeared in the expression-ranked heatmap, see colored names in Fig. 5F) and used them to color distinct regions of the spinal cord, highlighting the ability of this method to quickly organize transcriptomic “low-level” data into functionally significant “high-level” categories.

### 2.6 Regional and Cellular Morphology using Top-Down Analysis

With automated segmentation and region assignments, experimental questions can be asked directly from the segmentations. We used a publicly available Xenium dataset with four coronal hemi-sections of mouse brain. The set included coronal slices from two transgenic CRND8 (8 and 18 months) mice which are a model of Alzheimer’s disease and two WT (8 and 13 months) **(Fig. 6A**). We ran SiDoLa-NS-Macro-mCB on all the sections to extract the morphological data on single cells and atlas-defined brain regions (all in one step once parameters are set). We expected to see differences as the mice matured while also seeing small differences in brain regions in the older transgenic mice^72,73^. Even though neither younger mice nor Alzheimer’s samples were considered when generating the simulation dataset, the reference atlas was linked to each cell (**Fig. 6B**). While a single mouse per time point and condition is underpowered to ask biological questions, here we use it to illustrate the potential for cell and regional analyses on tissue. We first asked how the number of cells per region varied, and found the highest significance in the basolateral amygdala, the dentate gyrus granule cell layer, and the lateral amygdalar nucleus (**Fig. 6C** upper, F-stat 20.1, 61.4, 0.63). These all showed the same pattern of increasing cells with age in the WT but decreasing cells with age in the mutant. When cellular and regional inferences are performed, the neural network reports not only the predicted region label, but also its confidence in that call. Another interesting finding was that the confidence in calling many of the areas in the older Alzheimer’s brain diminished. This was especially notable in the somatosensory isocortex, in the auditory areas, and in the pallidum of this mouse (**Fig. 6C** lower, F-stat 6.6, 47.3, 11.85). The lower confidence could indicate that these areas are divergent, a possibility in line with the decline in mature CRND8 transgenic mice.

**Figure 6.**
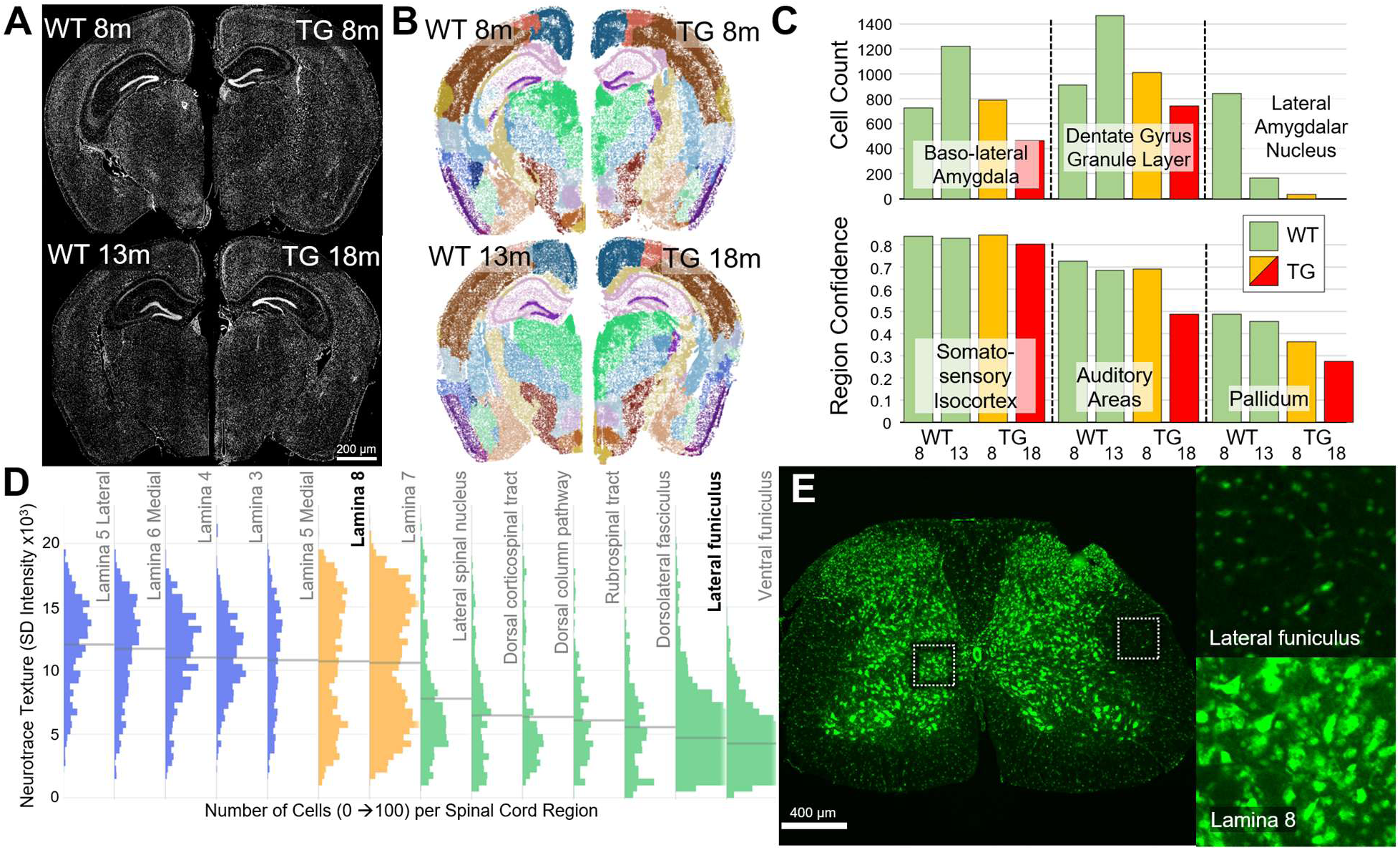
Utilization of Image Features to study Anatomical Regions in Mice. **A)** DAPI-stained mouse coronal hemi-sections from two wildtype (WT) mice at 8 and 13 months (left side), and transgenic CRND8 Alzheimer’s (TG) mice (right side). **B)** Diagram representing the same tissue sections but with markers representing single cells, and the coloration showing atlas regions, as defined by SiDoLa-NS-Macro-mCB (coloration matches Figure 2G). **C)** Bar charts measuring the number of cells per region (top), and the macro confidence within the region (bottom) in the four samples, WT 8 and 13 months, and TG 8 and 18 months. The bars are grouped by region, with Basolateral amygdala, dentate gyrus granule cell layer, and lateral amygdalar nucleus on the top, and Somatosensory Isocortex, auditory areas of the Isocortex, and pallidum below. **D)** Histograms showing the distributions of NeuroTrace texture (measured as standard deviation intensity) per cell, then separated by Spinal Cord regions. Horizontal lines show the mean for that region. Regions with low cell numbers were excluded. **E)** Micrograph of Spinal cord stained with NeuroTrace, where insets show a region of white matter (lateral funiculus) compared with Lamina 8, each with different NeuroTrace textures.

Next, we asked whether we could use morphological differences at the level of a cell (micro-scale), to find differences in the atlas regions within the mouse spinal cord section. The SiDoLa-NS inference automatically measures per-cell features for each available fluorescent channel. Indeed, we found that ‘NT cell texture’ (NeuroTrace intensity standard deviation amongst pixels per cell) arose, amongst others, as a strong segregator of spinal cord region (**Fig. 5D**, KW p∼0, H-Stat = 6139). In particular, *Lamina 8* and the *lateral funiculus* have distinctly different NT textures (**Fig. 5E**). Therefore, with only image-based information processed at two different scales, SiDoLa-NS-Micro-CNS and SiDoLa-NS-Macro-mSC enabled spinal cord region distinction utilizing subtle morphological features.

### 2.7 Simulation Engine for Multi Scale Nervous System Image Generation

The exclusive use of simulated microscopy images allows for generalizable identification of the nuclei or cell bodies, organized cellular clusters, and reference regions from anatomical atlases. However, these systems are only as capable as the simulated images from which they are trained. We asked whether we could develop a framework to produce *biophysically informed* simulations that would be compatible with the spatial omics platforms discussed in Figure 1. The pipeline below was used to create the simulations used in this manuscript but could also serve as a baseline from which to build other synthetic datasets for anatomical regions of interest.

Continuing the multiscale theme, this pipeline operates at micro-, meso-, and macro-scales (**Fig. 7A-C**). Each biological component is built as a 3D structure with a variety of properties defined and bounded by input parameters^51^. These components can be thought of as a function, and we represent them as a *node* with input parameters and outputs that return the geometry of that component (**Fig. 7D**). So, we first define the features of single cells (*Cell Node*), such as size, shape, and intensity, all of which can vary based on cell type and scan parameters and can be controlled through input parameters. The geometry for these (unique) individual cells is then used as an input to the next block (*Region Node*), which defines the makeup of a region or cluster (the details for the processes in the pipeline are defined in more detail below in Figure 6E and are indicated by a banner along the top). Cells are scattered throughout this region, with parameters for density and regional variation. The shape of these regions is either simulated generically, for example as laminar stripes, clusters, rings or is generated from a set of polygons which define a region from an anatomical atlas. Specifically, for sciatic nerve fascicles, we used rings of cells around a central ellipse of cells. For mesoscale inside the brain, we used higher density lamina that traverse across an image region as a stripe. Next, the geometry from each region is outputted and used to assemble the whole section (*Section Node*) that defines what will be a whole output image. Here, the parameters are randomized over a uniform random distribution to change the atlas morphology, so that it includes normal physiological variation but also goes beyond to encompass a variety of pathological phenotypes.

**Figure 7.**
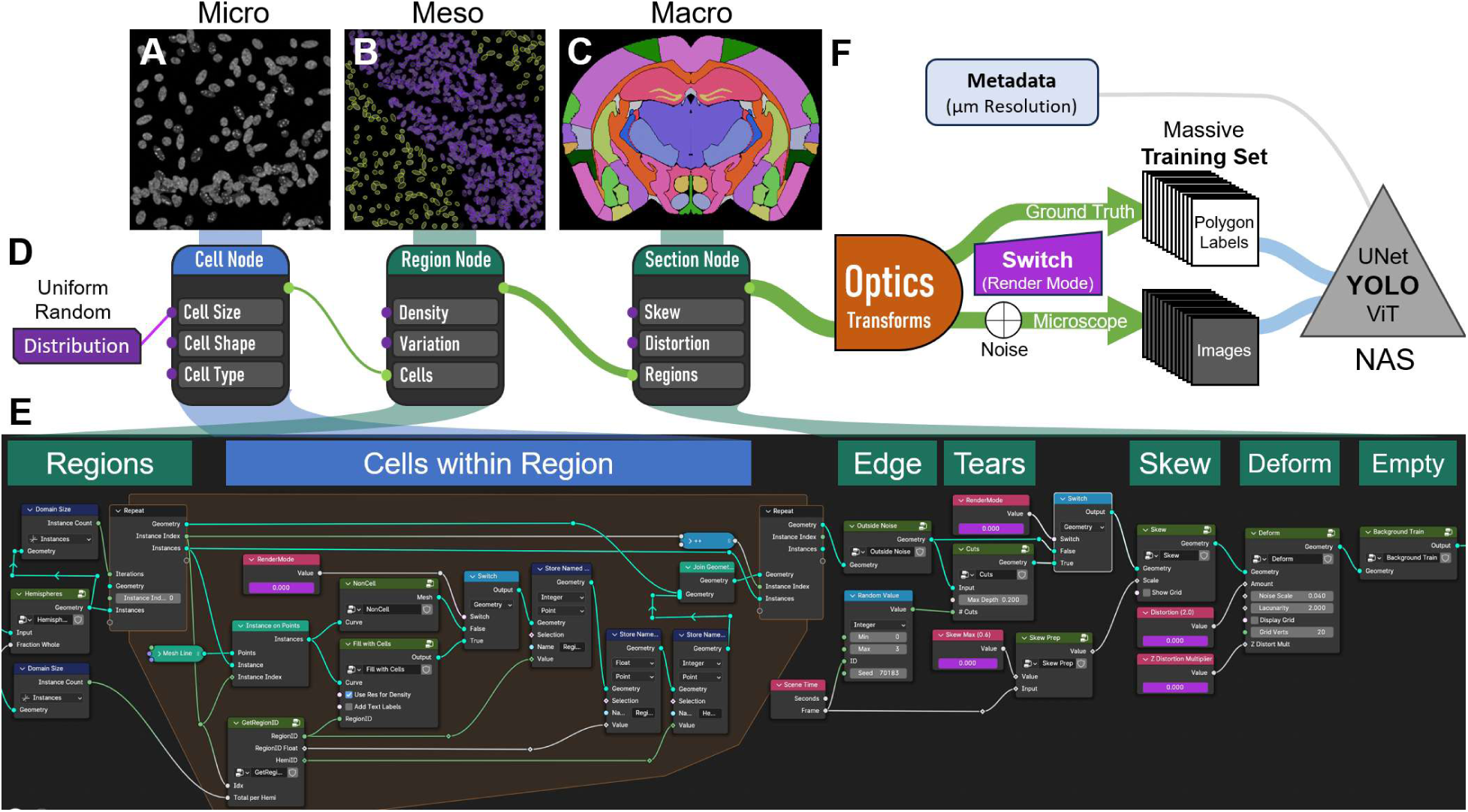
Simulation Engine for Creation of Synthetic Microscope Images and Corresponding Labels. The figure is organized to show the different modules used to produce the datasets for training, where **A-C** display micrographs and illustrations of the **A)** micro-, **B)** meso-, and **C)** macro-scale representations of, as an example, a coronal section of a mouse brain. **D)** Three “nodes” that represent the main concepts within the engine. These connect to the images above and the pipeline within Blender Geometry Nodes that execute each of the scales below. The simulation pipeline in **D** starts with a series of uniform random distributions, each of which control input parameters that define *Cell*, *Region*, and *Section* properties. The random seed is different for each distribution input, so that combinations of parameters are achieved. Within each node, several example parameters are listed, such as Cell *size, shape,* and *type*, with a circle on the left-hand side, indicating it is an *input* node. On the right-hand side, there is a green circle indicating an output node that passes 3D *geometry* on to the next node. The connections between the nodes increase in diameter, indicating that the geometry is increasingly complex as the smaller components are assembled into larger ones. **E**) The Geometry Nodes pipeline within Blender that produces the coronal brain and spinal cord simulations. Polygon regions that define a section from an atlas are the input, and then the central repeat zone (with a translucent orange background) iterates through each region. The center of this iteration defines the geometry of the individual cells that are placed therein. Then, section parameters are set, including *Edge*: placing debris and cells around the outside of the section, *Tears*: which simulates rips or tears, *Skew* and *Deform* which distort the section, and finally *Empty*: which occasionally includes empty/background scenes. **F**) The created geometry is run through an optic and physics engine resulting in an image. For each image set, two versions are created, a *ground truth* version and a *microscope* version. The switch that controls these changes parameters throughout the workflow, including the addition of noise at the end. The images are rendered along with corresponding ground truth polygons and labels. Metadata is also saved alongside the images that includes information about the pixel resolution and all the settings used. This dataset is then used to train a neural network, usually YOLO instance segmentation using neural architecture search (NAS).

We utilized the open-source CAD software Blender to produce the simulations, since it can be controlled through python scripting and with a visual generative workflow, called Blender Geometry Nodes (BGN), and has extensive optics and physics capabilities. The BGN pipeline for a micro-scale simulation focuses on cellular details (**Fig. S10**). A simplified version of the BGN pipeline that was used to create the macro mouse coronal brain dataset features transforms to distort the regions and overall section (**Fig. 7E**). The banner displayed over the pipeline defines the primary functions of each part of the pipeline: Regions, Cells within Region, Edge, Tears, Skew, Deform, and Empty. For the macro-scale reference regions, we defined their boundaries as exact copies of the reference polygons for one hemisection, and then had a switch to mirror those regions into a full set of hemispheres or just to use a single hemisection. The density of cells within the region was defined from a table (which we could manually set) or varied randomly between regions. Varying cell density randomly has the benefit of reducing our models’ dependence on density in learning region labels. Each region was produced iteratively, indicated under ‘Cells within Region’. The geometry from the regions and cells contained within were then joined together to define the whole section.

From here, the geometry of the simulated tissue section was manipulated. Specifically, we include within the BGN 1) Edge, which are extra cells and debris that are beyond the defined regions of the atlas (as there is often extra chunks of tissue that show up in a tissue section from errors in histological slicing), 2) Tears, which are meant to simulate rips or tears in the tissue. These two features are disabled when generating ground truth, as we want to reconstruct the tissue as if these imperfections didn’t exist. Additionally, we include two distortions, 3) Skew, which simulates different cut angles in sections and 4) Deform, which changes the relative size of the regions as would occur in samples from different individuals (the variation included purposely goes beyond common variation). Finally, there is a block that regulates 5) Empty, or background simulations, to reduce the influence of background variation and false positives in object recognition. More examples of these manipulations are shown in **Fig. S11, S12**. Although not shown in Figure 6, shading information is included to control how light is emitted from these objects and how it can interact with each object is also controlled using Blender Shader Nodes to enable variation and patterning.

At this point, the complex 3D geometry needs to be transformed into a 2D image, and the steps for that process are displayed below (**Fig. 7F**). To do this, we use optics and the rendering engine within Blender. Specifically, we employ a virtual camera with a 15.5 mm focal length and a low f-stop (0.01) to focus on the tissue section from about 1.4 mm away. Then the rendering engine ‘Cycles’ calculates the light being emitted by or bouncing off the samples and uses it to produce an image. After the image layer is produced, we use a compositor layer to add varying amounts of white noise (0-95%), making the image slightly harder to interpret. This process is repeated (per ‘frame’) for the total number of images and produces the simulated ‘microscope’ images for our dataset. At the same time, the software also calculates the boundary coordinates of the various objects within the scene from the camera’s point of view, and projects them onto a 2D plane so they are scaled to the rendered image. These coordinates are used to export the ground truth data, a set of polygon coordinates and class labels. For the ground truth construction, a switch is flipped that turns off some of the variation, such as the rips and the noise, so a clean reconstruction can be learned from the noisy microscope image. Also, during this process, all the parameters utilized are saved as metadata and are exported alongside the image and label files, so all the details can be tracked for different rendering runs and versions. In the macro mouse brain set, we rendered 4,065 images/hour on multiple desktops, with ∼700 image sets an hour on a computer with only an 8^th^ generation Intel CPU or NVIDIA 1000 series graphics card (i.e. standard, not specialty hardware).

The parameter input settings are constant for a single ‘frame’ of the BGN pipeline but are varied for different frames. Therefore, when producing the dataset, the software runs through one to hundreds of thousands of frames, creating the combinatorial variability of the training set. At the same time, we also produced a small, simulated validation dataset (hundred to thousand images). These images can then be fed into any number of image-based neural networks, such as a convolutional UNet architecture, instance segmentation architectures such as Mask-RCNN or YOLO^74^, or used with Vision Transformers (ViT). In this manuscript, we presented data from YOLO instance segmentation, and neural architecture search (NAS) was used to find the optimal settings. The metadata exported during rendering is loaded during training so that it can be later utilized during inference.

## 3 Discussion

Since Ibn al-Haytham’s pioneering work, optics has fascinated scientists. The microscopic world offers abundant data in the form of histological images—whether omics data is attached, these images form a valuable dataset in their own right. Even at the smallest scales, a single image of cells can spark whole fields of biology. Images of tissues are even richer, and some of the most complicated include the nervous system, where each subtly distinct region enables specific functions^75–77^. The complexity arises fundamentally from the tissues’ intricate structure, highly organized in the brain, spinal cord, nerves and support structures^78,79^. Understanding the nervous system in health and disease requires an in-depth exploration of the cellular and molecular patterns within their spatial context^80–83^.

Today, images carry vast additional dimensions, and hidden within vast pixel counts lie coupled transcriptomic and metabolomic data. For centuries, expert pathologists and histologists like Virchow, Malpighi, and Jacobi have discovered the embedded features in these datasets. As new technology promises to unveil the molecular underpinnings of tissue, it is likely that future tools will encompass all types of multi-domain omics and analyses^84,85^. This methodology aims to take a step in this direction by sharing a top-down approach to analysis that can be used in conjunction with spatial omics techniques.

Top-down simply means overlaying information that is directly extracted from the image itself, as opposed to building up that knowledge from a lower molecular level (bottom-up). SiDoLa-NS is built upon the premise that training a neural network using only available real images is insufficient. The variation in images that have been labeled is great, but the variation of a new section that needs to be analyzed is even greater. As with other advances in machine learning, this challenge can only be solved by a neural network when it has the chance to observe vast amounts of examples and a high degree of variation. Therefore, this technique uses “**Si**mulate, **Do**n’t **La**bel”.

The vanguard SiDoLa-NS models that we create with this approach are used in three distinct neurological cases, all with publicly available data. First, we infer cells, lamina, and atlas regions in a coronal slice of an adult mouse brain. While this works well, there are other approaches in the literature which can accomplish this task^44,86^. Next, we transitioned to two unique applications: a mouse spinal cord was segmented into its individual cells, and its reference atlas overlaid, identifying over 20 unique and identifiable regions accurately. In addition, we ran SiDoLa-NS on a cross section of sciatic nerve, which has distinctive morphology, including bundles of axons and Schwann cell nuclei organized into fascicles. Mice only have one or two of these fascicles, so we used a publicly available porcine section as a closer approximation of a human nerve. We demonstrate that these datasets can be used to identify marker genes when coupled with spatial transcriptomics. Additionally, we can use the multi-scale data to ask questions like how spatial distribution of neuronal cells in micro-scale correlate to the organization of spatial sub-regions of the corresponding tissue. Finally, we lay out the simulation process so that others can take advantage of the pipeline for their own applications.

The potential of SiDoLa-NS and other similar approaches is that it allows classifiers to become more and more general. Using NNs outside of the context they were trained in is fraught with problems, due to hallucinations and overfitting. One reason is that NNs can ‘memorize’ their training set, and another is variable performance on real samples because of batch effects introduced by the training set. Since we can simulate above and beyond the range of variation seen in existing samples, the neural network is more adept at handling variation in a completely new context. In fact, the NNs in this manuscript are used exclusively out-of-context since the training and validation data is simulated. Another subtle but important note is that the simulated training data is generated using an engine that is completely independent from training and validation. It does not use a GAN (generative adversarial network) or a diffusion approach to create the training set, which can lead to recursive generated collapse^87^. Other advantages of this approach include the generation of massive scales of training data, and the ability to overrepresent difficult and rare cases. For example, much of the mouse brain has nuclei spaced well apart, so existing cell segmentation NNs can handle these regions easily. But some regions, like the granule cell layers have extremely high cell density, producing consistent failures. With simulations, we can overrepresent these phenotypes, making the SiDoLa-NS models robust to high cell density when appropriately large bounds in image parameters are set (**Fig. S2**). Another important advantage of this approach is eliminating annotator bias, and that the quality of the ground truth annotations is much higher. For example, in the task of segmenting, the boundaries are now perfectly defined, since they are created from the simulation system at any resolution and the object bounds are defined before optical aberrations and sample imperfections, whereas manual annotations suffer from the vagaries of diffuse boundaries. Finally, since the simulation engine is based on biophysical properties (such as cells, clusters, and anatomically defined regions), its results are instantly and highly interpretable.

With these spatial-omics datasets, beware of double-dipping, which refers to using data in a circular way, such as identifying region based on molecular markers and then analyzing the same molecular signals within those regions. This approach compounds assumptions from both steps, leading to falsely inflated statistical significance. Although it is useful to enhance top-down image analysis with bottom-up clusters, the statistical significance of the results may be falsely inflated. Using top-down regional definitions sidesteps these problems^88–90^.

The power of SiDoLa-NS lies in the generation of the datasets. We chose to use those datasets with specific types of neural networks, but it is likely that additional exploration in that space would lead to even better inference. The NNs used in the manuscript are *instance* segmentation and/or *instance* classification models that are built on the YOLO^74^ Ultralytics framework. Instance segmentation is powerful since it does not just find the region, but it understands the difference between different regions and their identities. We also worked with MaskR-CNN^91^ and Feature Pyramid networks^92^ but found YOLO to have the most active development and efficiency. Still, YOLO and instance segmentation may not be the right approach for all applications. For example, anatomical region definitions involve non-overlapping areas, which is not an architectural constraint for instance-based methods. Instead, convolutional UNets perform better at these tasks, since each pixel’s location and identity are more consistent. In fact, we found that for some of these applications, a two-stage approach can enhance performance. First, UNets are trained to ‘recode’ the raw image with pixel-based classifications. Then, an instance segmentation model is trained from those, leading to better performance (data not shown).

We also have several improvements that are actively in development. During inference, the user must define the scale for which the SiDoLa-NS models should be run on the image. The next version of the inference system automatically checks the pixel resolution required by the model and lines it up with the input image. Also, in terms of scale, additional simulations are being produced at varying focal distances to improve performance with other microscope objectives or starting image resolutions. One major shortcoming of the current version is that the macro models are trained on only a single section from the reference atlas. While it is trivial to add additional sections into the training set, the next version will move beyond that. We are assembling all the slices in the atlas and using a system for virtual reconstruction into three-dimensional geometry^63^. Then, we can slice the atlas regions at any position, and at any angle. This will drastically improve the ability in multiple anatomical regions, in addition to helping deal with common histological issues like asymmetries in microtome slices. The current version has been extensively trained on both immunohistochemistry-like Nissl images, and immunofluorescence-like DAPI images. Although these current models can segment H&E, we plan to explicitly include these stains in training. The mouse brain, spinal cord, and peripheral nerves were chosen as important nervous system regions that could benefit from SiDoLa-NS. The upcoming versions will include additional nervous system tissue, including transverse sciatic nerve sections and dorsal root ganglia^93^ (both at micro- and meso-scales).

## 4 Methods

### Atlas Mapping

Resources from the Allen Software Development Kit (SDK) were utilized for the atlas mapping pipeline of the mouse brain and spinal cord. Each atlas image had a respective structure image with color-coded regions. The structure image for each atlas image was retrieved and a dictionary for regions and their respective color code was constructed. Next, binary masks in each color code were constructed over all structure maps to visualize the unique regions. Polygon coordinates defining the contours of each region were produced and assembled into a JSON file, organized by slice number and region name. These region outlines were used to generate macro-level brain and spinal cord simulations.

### Simulations Overview

All simulations were done in the CAD software Blender 4.0+ using a feature called ‘Geometry Nodes’ which allowed for a visual workflow that encoded the geometry representing different scales of biology. Individual ‘frames’ were used to control variations and produce the different training and test image datasets needed for teaching the neural networks. While each tissue type and scale used distinct geometries, a few general rules were common: The basic concept was to build a plane or volume to fill with cells, then to ‘instance’ cell objects and randomly fill them into the region. Both the regions, the fill density/pattern, and the cells were all created dynamically, and random variations were added for each frame. The more variation that was shown to the neural network, the better it was able to generalize on real tissue.

### SiDoLa-NS-Macro-CNS Mouse Coronal Brain and Spinal Cord

These sections were both produced with the same pipeline explained in Figure 6. After atlas mapping, the regions were imported into Blender as curve objects using a custom script. These were fed into the Geometry Nodes pipeline described extensively in Figure 6E. Additional details and images are found in Figure S10 and Figure S11.

### SiDoLa-NS-Meso-pSN Porcine Sciatic Nerve

Sciatic nerves were digitally generated using Blender with geometry nodes to simulate biophysical properties. The aim was to replicate natural structures within a sciatic nerve, including fascicles and Schwann cells. The Blender simulations were simplified to basic geometric forms, including ovular fascicles, circular Schwann cell nuclei, and circular epineurium. Additionally, varying sizes and counts of fascicles were distributed to ensure a manageable, yet flexible, simulation for training. This effectively prepared the model for real-world data. These simulated components were meticulously rendered into a set of .bmp files through Blender scripting, which produced the dataset.

### SiDoLa-NS-Micro-CNS Mouse Coronal Brain and Spinal Cord

The basis of this simulation was to instance ellipsoid nuclei on many points across a plane. The nuclei varied in size, aspect ratio, rotation, and shading (which featured bright heterochromatin spots). We also introduced two other meso-scale features. The first of which was a bright laminar stripe or striation. These were bands of much higher density nuclei compared to the adjacent cells (**Fig. S10**). The other feature was a region devoid of cells, usually thinner than the high-density lamina, that also crossed the image. Finally, a slight amount of depth-of-field with a low f-Stop was used to simulate slight changes in focus.

### SiDoLa-NS-Micro-PNS Mouse Peripheral Nervous System

This micro model is the original from which others were derived. It consisted of a plane that was filled with points that were split between two different cell type instances, each of which could be scattered in different ratios. One cell type was Schwann cell nuclei, which are more round, small and bright. The other type was the axons, which are slightly larger, often oblong, and dim to near invisible (by DAPI staining). The plane which contained these was circular shaped and had additional geometry outside of it representing the epineurium. The plane was also shifted around the frame of the camera so that occasional different edges of the perineurium were exposed.

### Optics, Noise, and Ground Truth Polygons

A virtual camera within Blender was used to translate the three-dimensional geometry into an image. This was done with the rendering engine “cycles” or “EEVEE” and involved calculating where the emitted light from the fluorescent objects fell or where light from the background in an IHC setting was blocked by the objects. The camera had depth-of-field and a small aperture to give slight out-of-focus regions. The camera was also tilted and twisted to give different viewing angles and raised or lowered to adjust the overall field-of-view and zoom. White noise was added on top of the final image with the compositor and mixed at different ratios to make the SiDoLa-NS models robust against noise when inferring. In addition to the standard rendering, we also extracted the coordinates of the vertices making up the 3D geometry, and projected them onto a plane from the camera’s point of view, converting them into camera x,y coordinates (world to camera). These vertices were then reassembled into polygons directly, or convex hull was applied. In more complex setups (atlas regions), we also down sampled the resolution of the polygons so that the exported segmentation files were more manageable. These files were jointly exported in Yolo.txt or coco.json format automatically as the images were rendered.

### Model Training

The images created could be used with a variety of instance segmentation or object detection architectures. All the inferences in this document utilized Ultralytics YOLO v8 or v9 pre-trained foundation models in a python environment with CPU or GPU. For micro models, where the exact outline of the nucleus was not required, we trained ellipsoid nuclei segmentations into bounding boxes with YOLO v9c and v9e. For more complex shapes, like in the macro models, we trained with YOLO v8n-seg and b8l-seg foundation models to extract the full polygon masks from the inferences. The number of images used in training is listed throughout the results. Generally, the training was for 10-200 epochs, the image size was between 448 and 640 pixels in width and height, and the batch size was between 8 and 32 images per batch. All these factors were varied, along with the random seed, for each NAS run.

The results of the run were copied into a folder that was named by the resulting mAP_75_ on the validation dataset, and then inferred on biological examples as a quick reference.

### Transcript Processing

The 10X Visium HD Brain, acquired from a public source, was provided at 2, 8, and 16 µm resolution in the binned outputs folder. We used pandas to export the tissue parquet files as CSV files, acquiring the SRT data in micron coordinates. The 10X Xenium Spinal Cord was provided as a CSV file, with individual transcripts identified by micron coordinates. The CSV mapped directly to the DAPI-stain image of the tissue sample. Given the slice’s width and height in microns, SiDoLa-NS automated the scaling required to map the transcripts to the segmented cells.

### Data Collection and Image Preprocessing

#### Mouse Coronal Brain

The Visium HD Brain H&E image was acquired from 10X Genomics. The microscopic image provided was an H&E-stained section. Utilizing ImageJ, we split these images into its constituent channels and selected the channel that best highlighted nuclear features (red channel) for further processing. For both cases, we adjusted the image brightness and contrast, applying a linear normalization to the whole image. For macro/meso/micro inference, these images were directly given to the program, without down sampling or tiling.

#### Mouse Alzheimer’s Brains

Four brain hemi-sections from a publicly-available dataset, “Xenium In Situ Analysis of Alzheimer’s Disease Mouse Model Brain Coronal Sections from One Hemisphere Over a Time Course - 10x Genomics” were downloaded. The DAPI images were rotated to be upright in ImageJ, and then targeted for a macro/meso/micro inference as above. Statistics for effects on these tissues were linear regression against this scale: WT 8 = 0, TG 8 = 0.2, WT 13 = 0, TG 18 = 1 for regional confidence, and this scale WT 8 = 1, TG 8 = 1, WT 13 = 1.5, TG 18 = 0.5 for cell numbers per region.

#### Mouse Spinal Cord Immunofluorescence

For the spinal cord analysis, we acquired the IF images from a spinal atlas manuscript^64^. The spinal cord dataset included nine different cross sections of spinal cords, each containing three different channels of ChAT, Nissl, and DAPI stains. We selected *Slide1-6_Region0007_Channel555 nm,475 nm,395 nm_Seq0051* and labeled each spinal cord from one to nine starting from the top and moving horizontally. Taking the second spinal cord, we cropped it into a square and split it into three separate channels using ImageJ. The Nissl channel was then used for our SiDoLa-NS models’ predictions.

#### Mouse Spinal Cord Xenium

A C57Bl6-J female, 109 days old mouse was anesthetized in an isoflurane induction chamber perfused sequentially with ice-cold RNase-free PBS followed by fresh 4% formaldehyde (Sigma Aldrich, cat. 100496). To preserve RNA integrity, the entire spine, including the spinal cord, surrounding bone, and musculature, was rapidly dissected and incubated in 4% formaldehyde at 4°C for 48 hours. After fixation, the surrounding bone and muscle structures were removed by dissection, and the exposed spinal cord was transferred to 70% ethanol at 4°C until processing. The fixed spinal cord tissue was dehydrated and embedded in paraffin using a Tissue Processor (Leica TP 1020) through graded ethanol, xylene, and melted paraffin immersions. The spinal cord sample was then embedded in a cold paraffin block, sectioned into 6 µm slices with a microtome, and floated in a 42°C water bath. Tissue sections were mounted on Xenium slides (10X Genomics, PN-1000460) within the 12 mm x 24 mm imageable area. Slides were dried for 30 minutes at room temperature, incubated for 3 hours at 42°C in a dryer oven, and placed in a desiccator to dry overnight at room temperature. Finally, the slides with the lumbar region were hybridized with 10X Mouse Brain Panel probes (which included ChAT) and processed using the Xenium Analyzer following the manufacturer’s instructions.

#### Porcine Sciatic Nerve

The porcine sciatic nerve slide was obtained from Saarland University’s histology site https://mikroskopie-uds.de/^94^. The sciatic nerve cross section had a scan area of 6.2 mm x 4.8 mm and H&E staining. The image was downloaded as a .zif file, which was then loaded into ImageJ and the highest resolution version of the sciatic nerve was exported as a 38885 x 30286 pixel 8-bit image. The channels were then split. Both channel 1 and channel 2 highlighted the fascicles within the sciatic nerve. Due to channel 2’s more distinct contrast between the fascicles and the background, channel 2 was used for Figure 4. Channel 3 highlighted the axons within the fascicles. The pixels were then inverted to create a black background with white fascicles and adjusted to enhance the visibility of the fascicles and axons against the background and to mimic DAPI staining.

### Inference

SiDoLa-NS must work on all parts of the image at different scales, and then that information needed to be fused together, along with any other omics data that corresponded with the image dataset. This was accomplished with the macro/micro inference package included in the code repository. The input allowed a single micro-scale model to do the cell segmentation, and then allowed as many meso- or macro-scale models to be included. Some parameters were set up for each model such as the location of the images, and then the software ran all the inferences and linked the data together. For the micro-scale, the multiple channels available from the image were measured across three masks: Box, Poly, and Voronoi (which represent the bounding box, the full polygon outline if available, and the space all around the cell until the next cell’s boundary is reached). Area, Size, Intensity, and Texture (SD) were measured for each mask and reported on a per-cell basis. If there was more than one macro-scale model, then hard and soft-voting ensembles were generated and reported.

### Analysis

#### Model Evaluations

SiDoLa-NS models were evaluated during training utilizing the Ultralytics *validation* tools. For all our models, the validation set used during training consisted of simulated images. For each validation image, many metrics including precision, recall, IoU, and confidence were recorded. Precision and recall were used to generate an 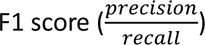 as well as graphed across confidence values. Additionally, a confusion matrix was generated for visualization of classification accuracy, particularly in multi-class tasks like brain and SC atlas mapping.

#### Robustness Analysis

Various robustness analyses were conducted on the macro models for noise, degree of rotation, and focus. We generated a small set of 18 – 25 images with calibrated variations in one of the three ways. For noise, we altered the image resolution by reducing the pixel contrast. For degree of rotation and zoom, the camera was respectively rotated or moved along the z-axis relative to the reference object. We also adjusted the F-number for the camera’s aperture for focus. For each of those images, we additionally generated a ground truth, which was used in evaluation. The mAP_50_ was primarily utilized in assessing model performance.

#### Marker Gene Analysis

Because SiDoLa-NS-Macro-mCB and SiDoLa-NS-Micro-CNS were inferred directly on the VisiumHD mouse brain slice, we mapped the gene expression matrix at the 16 µm bin level to the data based on micron position. For the Xenium spinal cord slice, SiDoLa-NS-Macro-mSC and SiDoLa-NS-Micro-CNS were both evaluated, and transcripts automatically assigned to cells. This resulted in each detected cell having a brain region assignment and a known gene count. For the list of the ‘top ratio’ genes, we ran a Kruskal Wallis test including all genes against brain regions to determine the highest rank of genes. From there, the ratio of each gene’s count to the total region gene counts allowed us to rank these genes per region, yielding our list of the top 4 genes.

#### UMAPs and Clustering

The UMAPs were viewed using the Seurat embeddings from the VisiumHD dataset at the 16um bin level. The UMAP1 and UMAP2 data was plotted and colored by cluster. For the cluster definitions, we utilized the provided graph-based clusters, which we joined to the UMAP with the barcodes. The SiDoLa-NS UMAP in Figure 2G uses the same underlying projection data but was colored with the SiDoLa-NS-Macro-mCB predictions for the respective barcode region.

## Abbreviations

SiDoLa-NS: Simulate, Don’t Label – Nervous System
SiDoLa-NS-Micro-CNS: Micro – Central Nervous System
SiDoLa-NS-Micro-PNS: Micro – Peripheral Nervous System
SiDoLa-NS-Meso-pSN: Meso – porcine Sciatic Nerve
SiDoLa-NS-Macro-mCB: Macro – mouse Coronal Brain
SiDoLa-NS-Macro-mSC: Macro – mouse Spinal Cord
IoU: Intersection over union
mAP: Mean average precision
YOLO: You Only Look Once; image detection and segmentation foundation models

## Acknowledgements

Thanks to Jane Dodd at Columbia and Felix Fiederling for their paper providing the spinal cord images. Thank you to Rudolf Bock at Saarland University for their virtual microscope histology site that provided the porcine sciatic nerve cross section. Thanks to Ken Lau and Harsimran Kaur at Vanderbilt University School of Medicine for helping with their MILWRM code. We would also like to thank Serena Elia, Graham Bachman, Purva Patel, Jimin Lee, Josh Milbrandt, and Saul Weiss, for their help with aspects of the project. Additionally, we would like to thank the Milbrandt, DiAntonio and Mitra labs for their continued support. We would like to thank the **McDonnell Genome Institute**, and the rest of the FIVE@MGI lab members. Finally, we would like to thank the Genetics and MGI administration and our maintenance and cleaning staff.

## Author Contributions

L.K.MA., A.K.Y.Y., E.B.G., T.N., N.K.T., F.P.I., R.S., D.G., J.E.W., C.L.K., and W.J.B. wrote the manuscript. W.J.B., L.K.MA. designed the experiments. Initial planning and ideas brought by W.J.B., J.D.M., R.D.M., A.K.Y.Y., D.G. and L.K.MA. L.K.MA, A.K.Y.Y., T.N., N.K.T., W.D., and L.Y. performed the experiments. L.K.MA., A.K.Y.Y., E.B.G., T.N., N.K.T., F.P.I., R.S., and W.J.B. analyzed the results. Key Software was developed by W.J.B., L.K.MA., and N.K.T. All reviewed the paper and gave suggestions.

## Data and code availability

https://gitlab.com/buchserlab/sidola-ns

https://huggingface.co/collections/FIVE-MGI/sidola-ns-676d91b211b32e84adbf391f

## Notes

https://gitlab.com/buchserlab/sidola-ns/

https://huggingface.co/collections/FIVE-MGI/sidola-ns-676d91b211b32e84adbf391f

### Competing Interest Statement

The authors have declared no competing interest.

### Summary of Updates

Figure 5 and the corresponding section was split to new Fig 5/ fig 6. New supplemental figure added comparing to other methods.

