## Supplemental Figures for "Biophysical Simulation Enables Multi-Scale Segmentation and Atlas Mapping for Top-Down Spatial Omics of the Nervous System"

Alternative Title: Biophysically informed Nervous System Spatial Image Analysis at multiple Scales

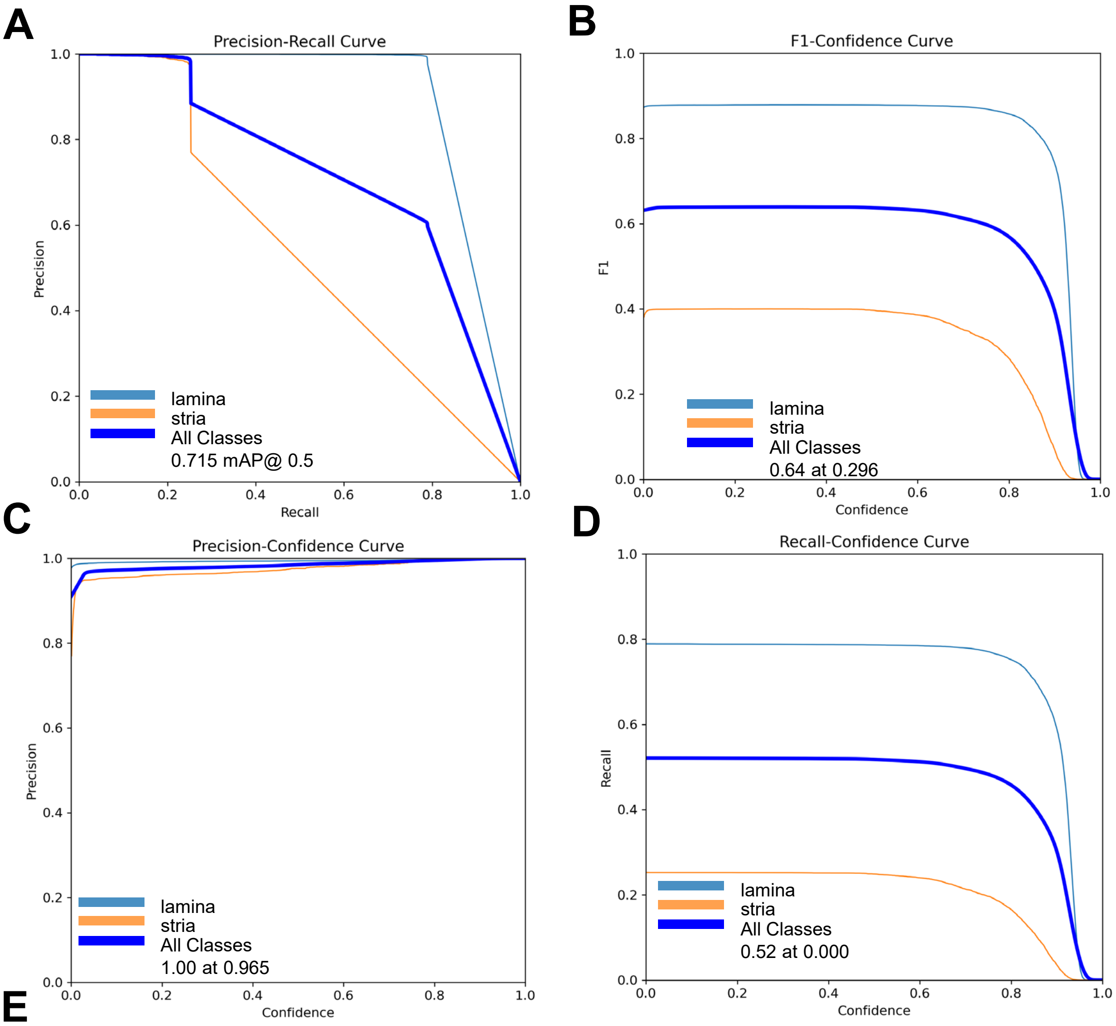


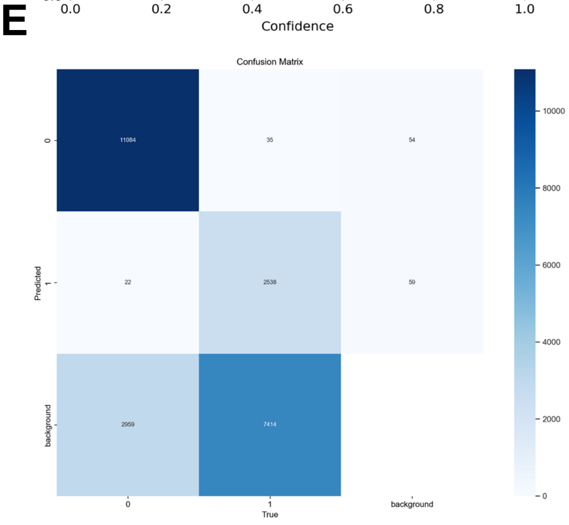


**Figure S1. SiDoLa-NS-micro-mCB model performance statistics.** For all panels, light blue curves represent 'lamina', orange curves represent 'stria', and dark blue curves ('All Classes') represent metrics averaged across all classes. **A.** Precision-Recall curve demonstrating how the model average precision for bounding box predictions with IOU > 0.5 varies as a function of recall. The max P for All Classes is 0.715 **B.** F1-Confidence curve showing how F1 score for bounding box predictions varies as a function of confidence. The top F1 score for All Classes is 0.64 and is reached at a confidence of 0.296. **C.** Precision-Confidence curve showing how Precision for bounding box predictions varies as a function of confidence. The top P for All Classes is 1.00 and is reached at a confidence of 0.965. **D.** Recall-Confidence curve showing how Recall for bounding box predictions varies as a function of confidence. The top R for All Classes is 0.52 and is reached at a confidence of 0.000. **E.** Confusion matrix for the model class predictions. Values in the boxes are instance counts for each class in the predictions versus true labels. Darker boxes represent greater values.

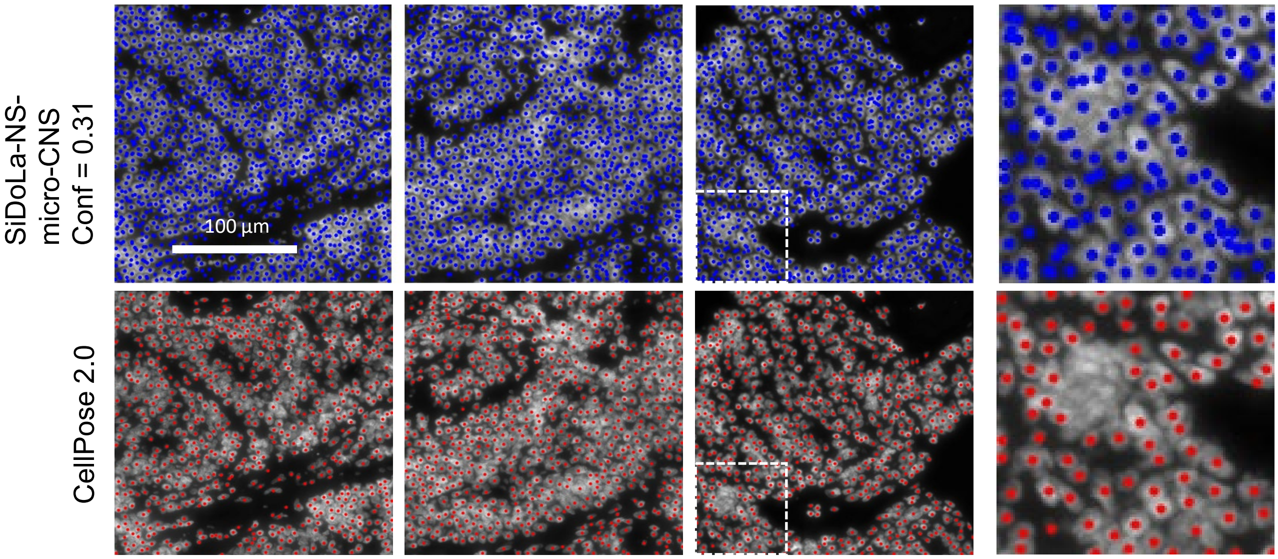


**Figure S2. SiDoLa-NS-micro-CNS performance in Lung Adenocarcinoma.** To test model performance on alternative challenging cases, SiDoLa-NS-micro-CNS was inferred on a DAPI-stained cross section of Lung Adenocarcinoma provided by 10x Genomics. The top row shows SiDoLa-NS-micro-CNS performance on a few high-density tiles. The bottom row shows the CellPose 2.0 prediction for the same images. Blue (SiDoLa-NS-micro-CNS) and red (CellPose 2.0) dots illustrate detected nuclei overlayed over the original images. A white box denotes the region corresponding to the zoomed image on the far right. White bar represents the image scale.

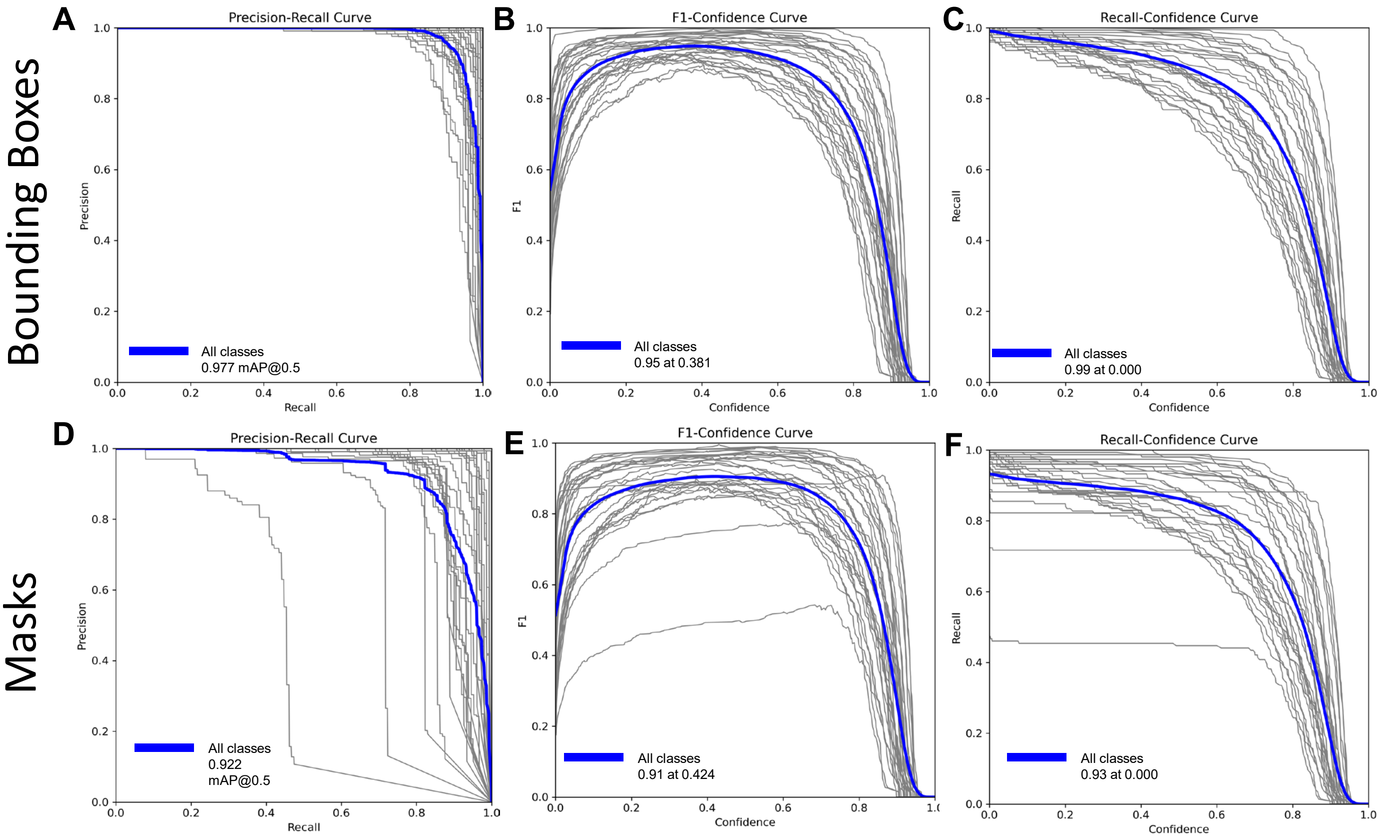


**Figure S3. SiDoLa-NS-macro-mCB model performance statistics.** For all panels, grey curves represent individual classes, and the blue curve ('All Classes') represents metrics averaged across all classes. **A.** Precision-Recall Curve demonstrating how the model average Precision for bounding box predictions with IOU > 0.5 varies as a function of recall. The max Precision for All Classes is 0.977 **B.** F1-confidence curve showing how F1 score for bounding box predictions varies as a function of Confidence. The top F1 score for All Classes is 0.95 and is reached at a confidence of 0.381. **C.** Recall-Confidence curve showing how model recall Recall (R) for bounding box predictions varies as a function of confidence. The top Recall (R) for All Classes is 0.99 and is reached at a confidence of 0.000. **D.** Precision-Recall curve for mask predictions with IOU > 0.5. The top P for All Classes is 0.922. **E.** F1-Confidence Curve for mask predictions. The top F1 score for All Classes is 0.91 at confidence of 0.424. **F**. R-confidence curve for mask predictions. The top R is 0.93 at confidence 0.000.

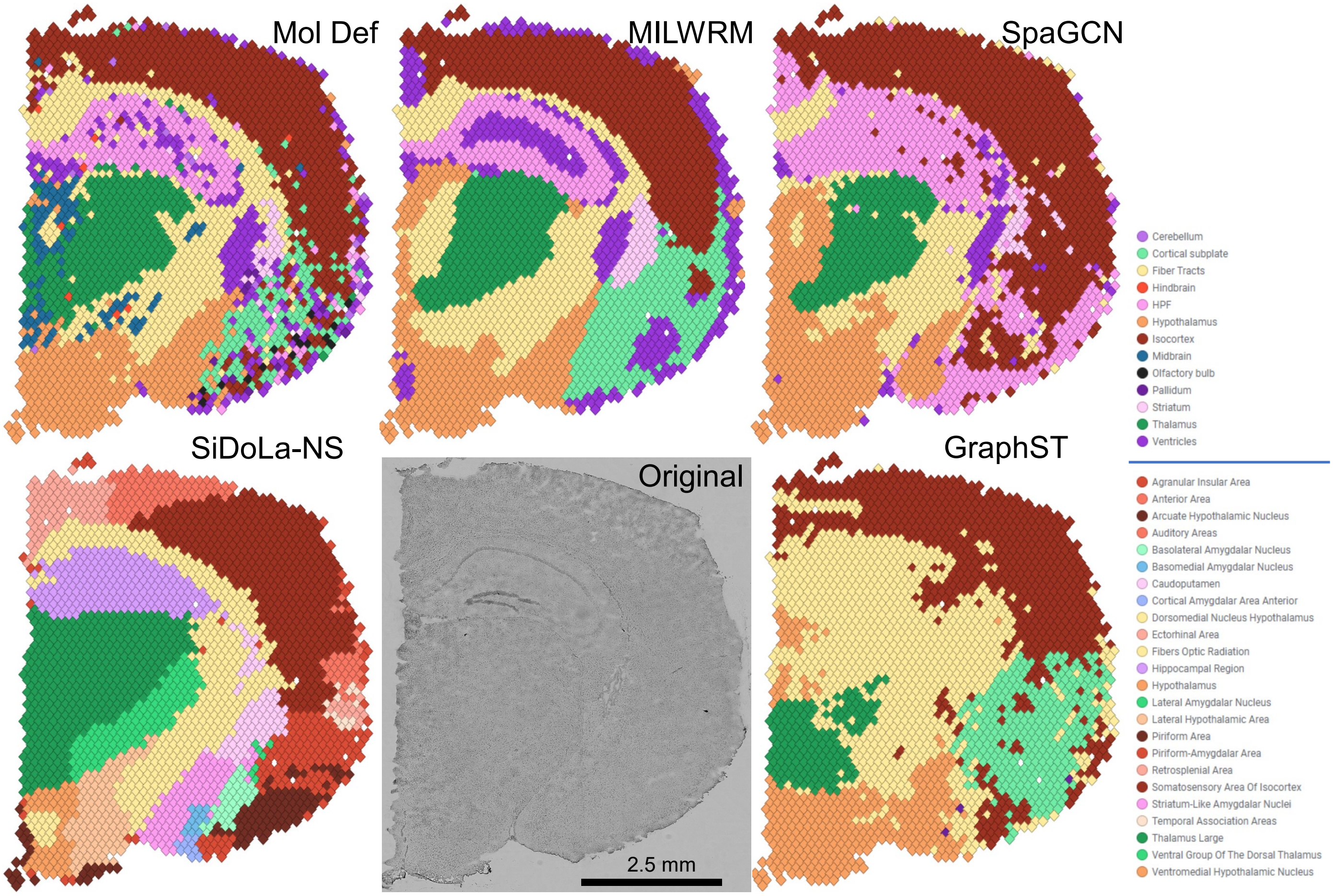


**Figure S4. SiDoLa-NS-Macro-mCB and comparisons to Expression-Defined Regions in Visium Standard Datasets.** Detection of tissue domains in a mouse-brain coronal section. Inference was performed on an H&E-stained coronal section of a mouse brain provided by 10X Genomics Visium Spatial Transcriptomics Dataset (termed Original). In the other graphs, each unique color (upper legend) corresponds to a distinct mouse brain region defined by expression patterns in MILWRM's manuscript (Mol Def). SiDoLa-NS has additional color variants to represent subregions (lower legend). The data from MILWRM's repository (https://github.com/Ken-Lau-Lab/MILWRM) was used to evaluate Mol Def, MILWRM, SpaGCN, and GraphST. We gave further advantage to the other techniques since their labels are simply numeric indices of clusters, and we assume that the cluster best overlapping with the molecular definition should get the molecular-defined label. It is also worth noting that none of these expression-based methods directly name regions and instead require manual assignment. We had to down-sample the SiDoLa-NS labels to fit within the smaller subset of the MILWRM molecular-defined labels (from 24 to 13) for comparison. We treated each method as observations and compared with the MILWRM Mol Def as ground truth to generate a confusion matrix and evaluation metrics.

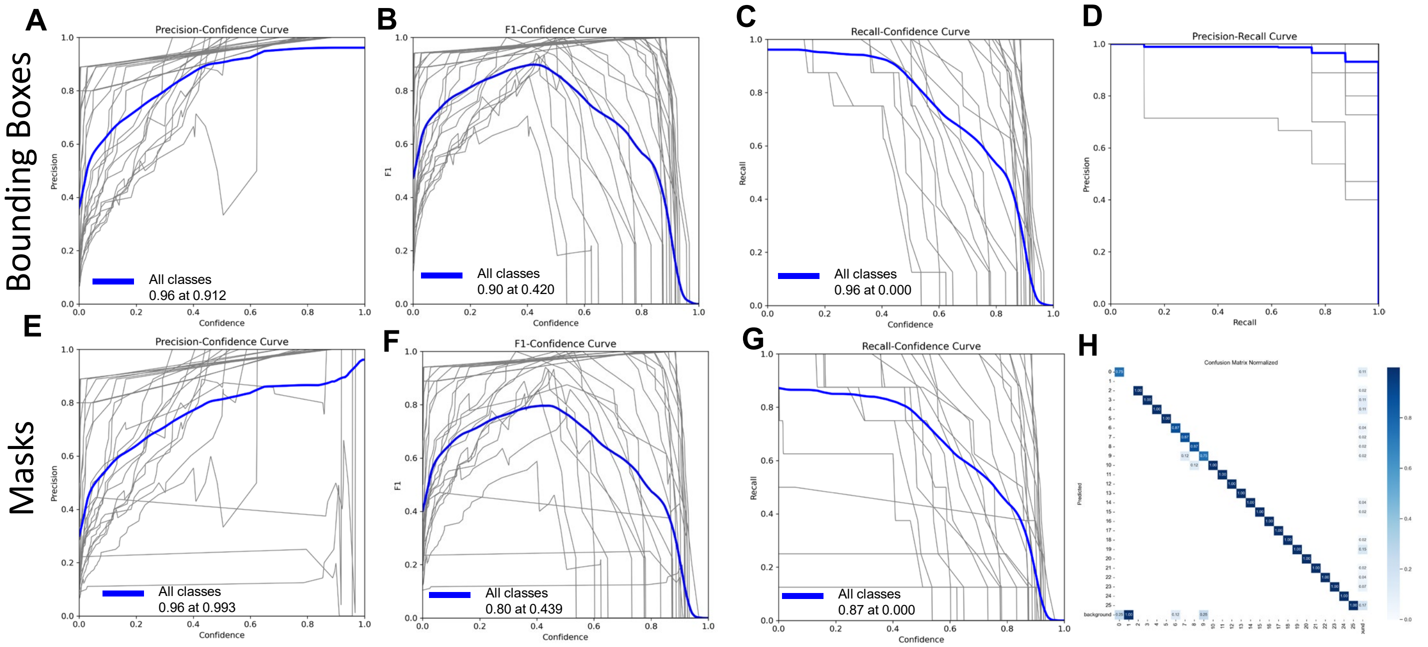


**Figure S5 Spinal Cord Macro Eval Metrics.** For all panels, grey curves represent individual classes, and the blue curve ('All Classes') represents metrics averaged across all classes. **A.** Precision-Confidence curve demonstrating how the model average precision (P) for bounding box predictions with IOU > 0.5 varies as a function of confidence. The max P for All Classes is 0.96 at 0.912 confidence **B.** F1-confidence curve showing how F1 score for bounding box predictions varies as a function of confidence. The top F1 score for All Classes is 0.90 and is reached at a confidence of 0.420. **C.** Recall-Confidence score showing how model recall (R) for bounding box predictions varies as a function of confidence. The top R for All Classes is 0.96 and is reached at a confidence of 0.000. **D.** P-R curve for mask predictions with IOU > 0.5. The top P for All Classes is 0.937. **E.** P-confidence curve for mask predictions where IOU > 0.5. The top P is 0.96 at confidence 0.933. **F.** F1-confidence curve for mask predictions. The top F1 score for All Classes is 0.80 at confidence of 0.439. **G**. R-confidence curve for mask predictions. The top R is 0.87 at confidence 0.000. **H.** Confusion matrix for the model class predictions. Values in the boxes are instance counts for each class in the predictions versus true labels. Darker boxes represent greater values.

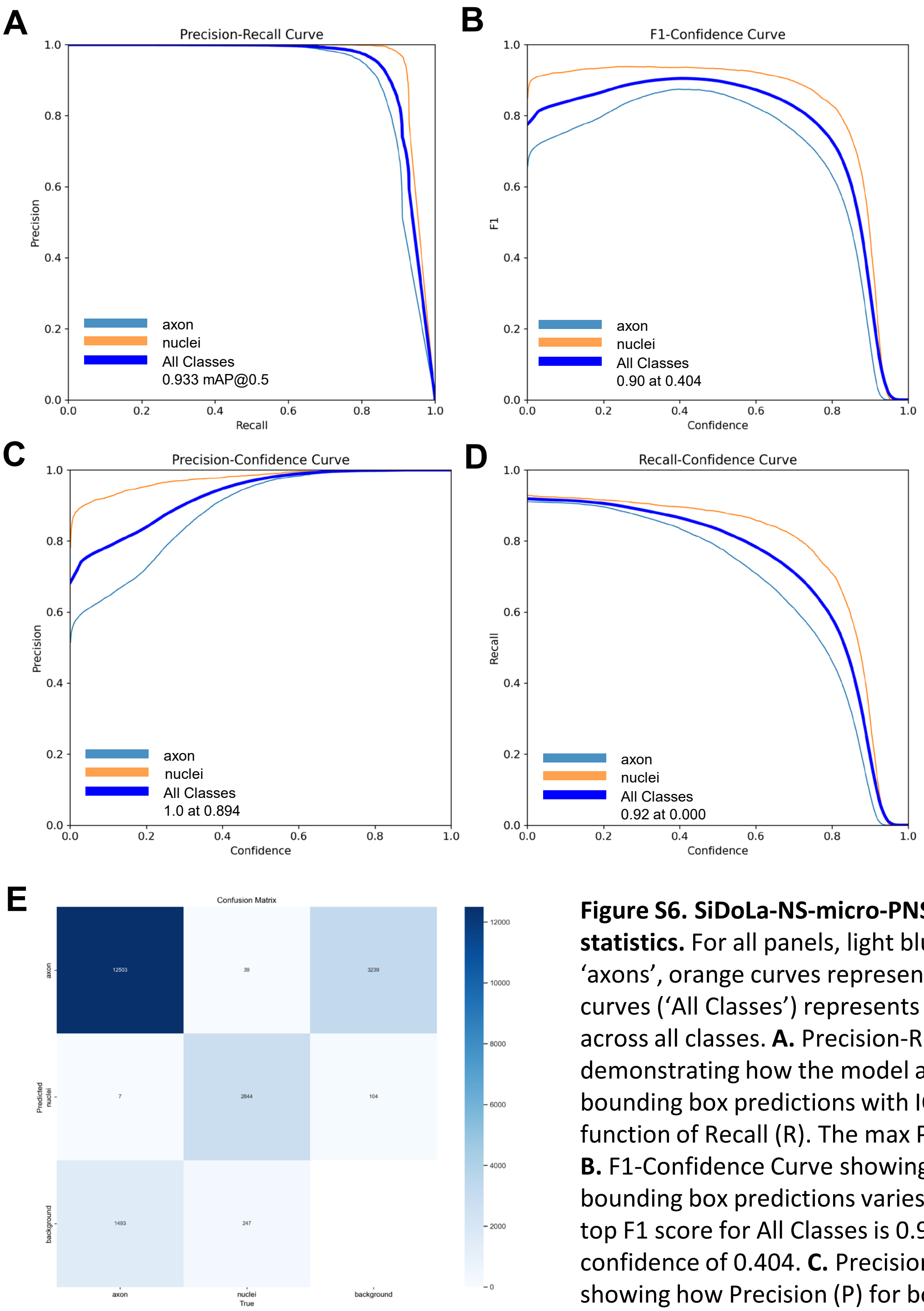


**Figure S6. SiDoLa-NS-micro-PNS model performance statistics.** For all panels, light blue curves represent 'axons', orange curves represent 'nuclei', and dark blue curves ('All Classes') represents metrics averaged across all classes. **A.** Precision-Recall Curve demonstrating how the model average Precision (P) for bounding box predictions with IOU > 0.5 varies as a function of Recall (R). The max P for All Classes is 0.933 **B.** F1-Confidence Curve showing how F1 score for bounding box predictions varies with confidence. The top F1 score for All Classes is 0.90 and is reached at a confidence of 0.404. **C.** Precision-Confidence Curve showing how Precision (P) for bounding box predictions varies as a function of confidence. The top P for All Classes is 1.00 and is reached at a confidence of 0.894. **D** Recall-Confidence Curve showing how Recall (R) for bounding box predictions varies as a function of Confidence (C). The end of the Recall plateau for All Classes is 0.91 and is reached at a confidence of 0.2. **E.** Confusion matrix for the model class predictions. Values in the boxes are instance counts for each class in the predictions versus true labels. Darker boxes represent greater values.

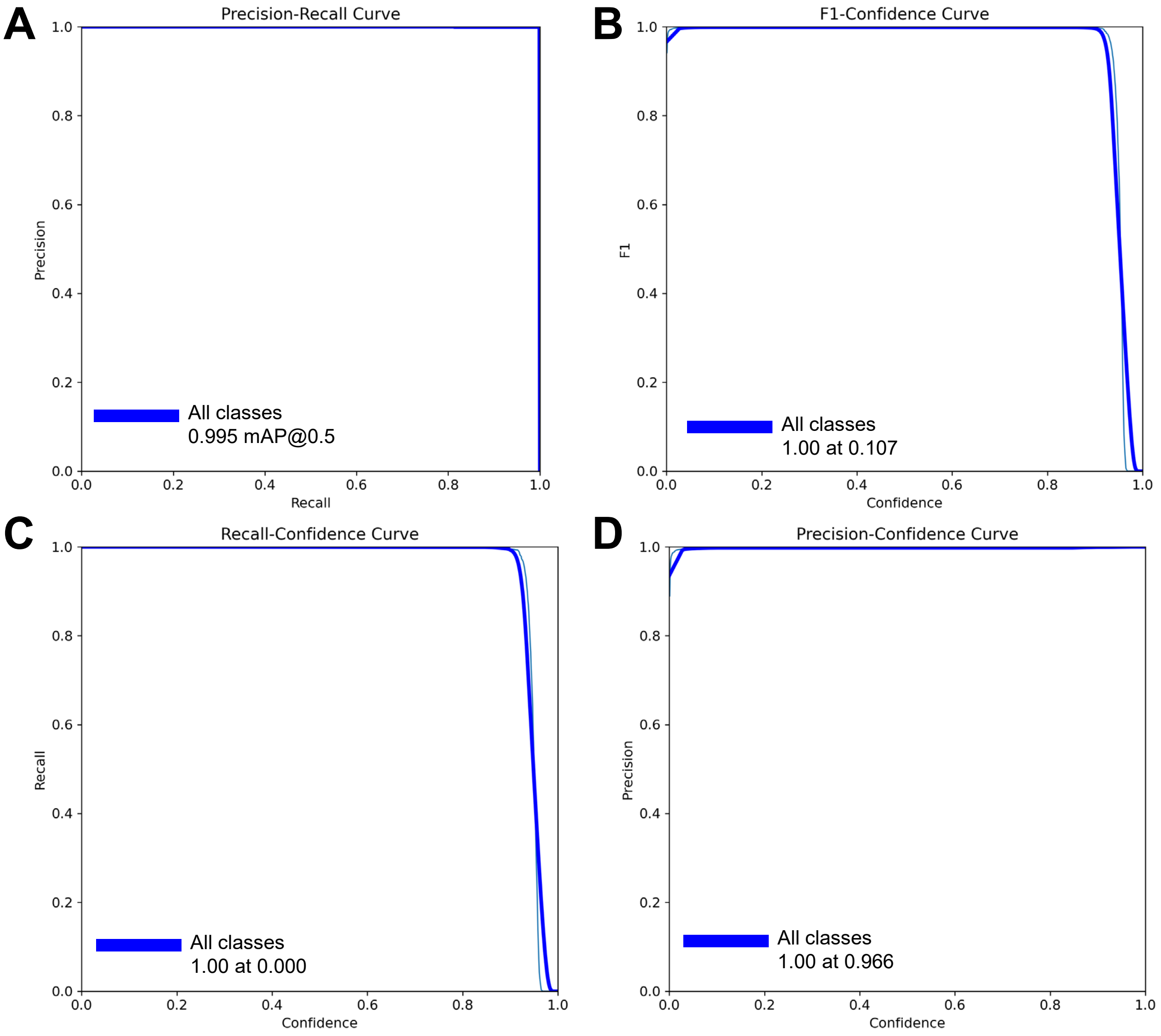


**Figure S7: Fascicle Meso Performance Metrics A.** Precision-Recall Curves for various classes are shown: Fascicles (light blue) and Fascicle + Background (blue). Mean average precision score is at a mAP 0.5. The top Precision for Fascicle + Background was 0.995. **B.** F1-Confidence Curves for various classes are shown: Fascicles (light blue) and all classes (blue). The curve demonstrates the trade-off between F1 and confidence at different classes, with the peak F1 score of 1.00 indicating the optimal confidence threshold is at 0.107. **C.** Recall-Confidence Curves for various classes are shown: Fascicles (light blue) and all classes (blue). The curve demonstrates the trade-off between recall and confidence at different classes, with the peak recall score of 1.00 indicating the optimal confidence threshold is at 0.000. **D.** Precision-Confidence curves for various classes are shown: Axon (light blue) and all classes (blue). The curve demonstrates the trade-off between precision and confidence at different classes, with the peak precision score of 1.00 indicating the optimal confidence threshold is at 0.966.

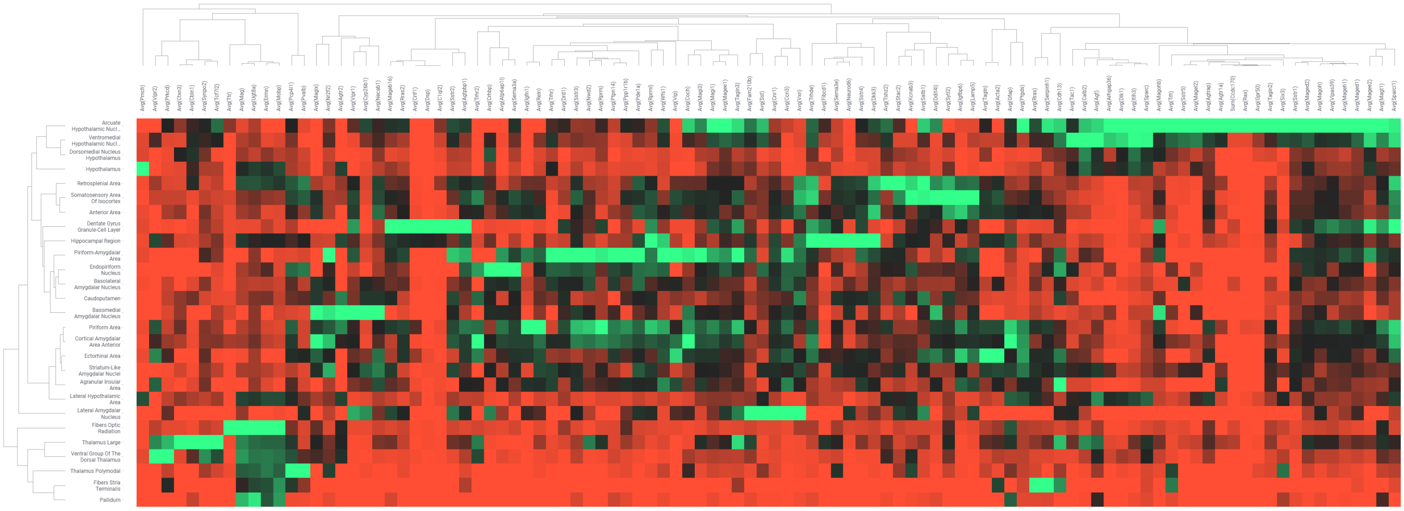


**Figure S8. Mouse Brain Marker and Regional List from Top-Down Analysis.** Heat-map represents the gene expression data for 102 of our top spatially variable genes. On the y-axis are brain region and on the x-axis are the genes. Both axes are grouped by similarity. Color represents average gene counts, where bright green and bright red are respectively the high and low maxima for gene count per region. Due to the large number of axes, it is difficult to discern a diagonal, although hotspot regions are visible.

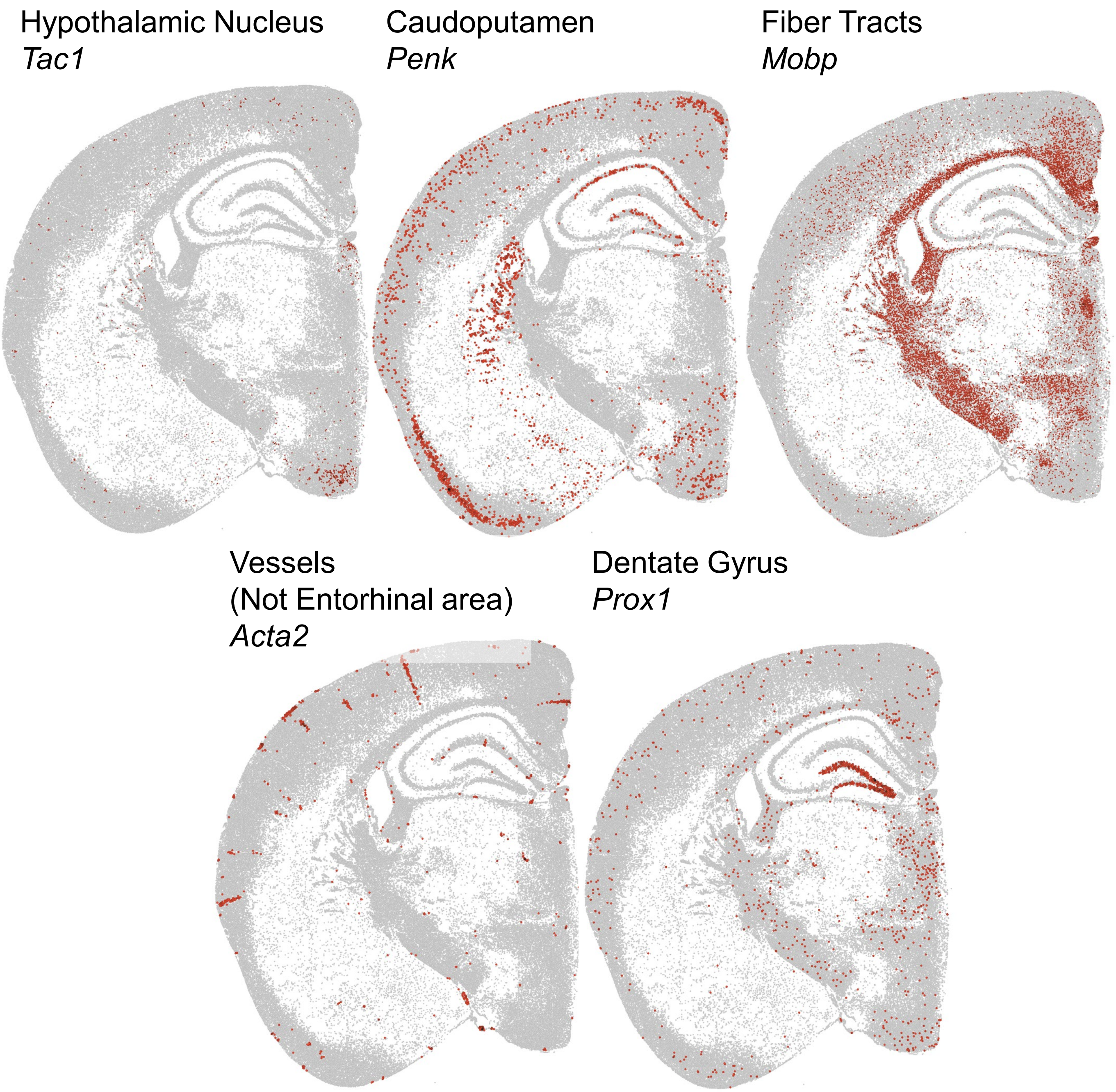


**Figure S9: Unique Marker Details.** Mouse brain, cell-based examples showing Sum Counts per Nuclei for Gene specified, where gray is 0, mean is black, and max is red. Five genes are highlighted (from left to right): *Tac1* in the hypothalamic Nucleus, *Penk* in the Caudoputamen, *Mobp* in the fiber tracts, *Acta2* in the Vessels, and *Prox1* in the dentate gyrus. *Acta2* is an interesting example of a gene which SiDoLa-NS-macro-mCB mis-assigned. While this was one of the top differentially expressed genes in the Entorhinal area, visualizing its expression reveals its restriction to the vessels.

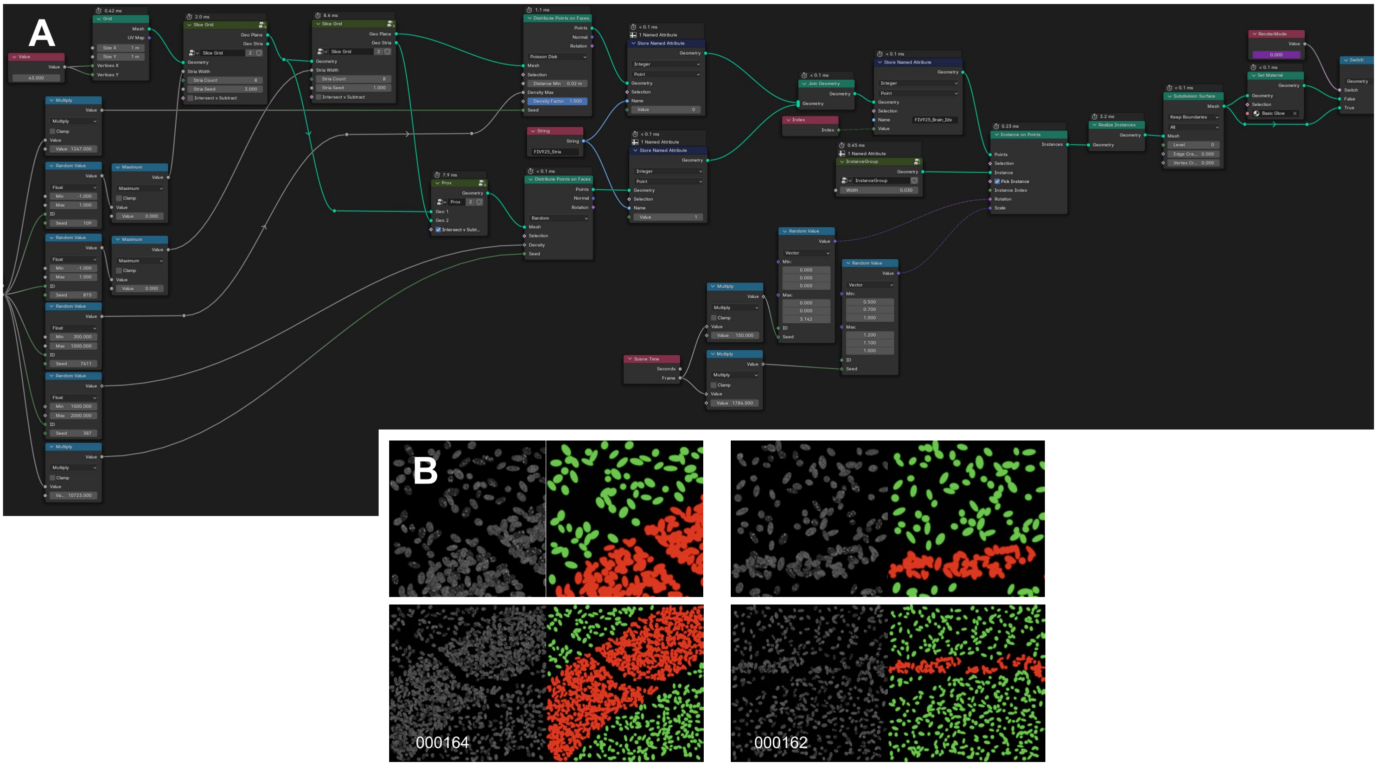


**Figure S10. Micro Model Simulation Examples. A.** Geometry nodes used to generate the micro- and meso-scale image sets of the central nervous system. **B.** Examples of rendered meso-scale images. In white text is the image's unique ID. Each colored image is the label for the respective image to its left. The bottom row is a more zoomed out version of the row above to better demonstrate the tissue striations. Green represents adjacent cells; red represents high density stria or bands of cells.

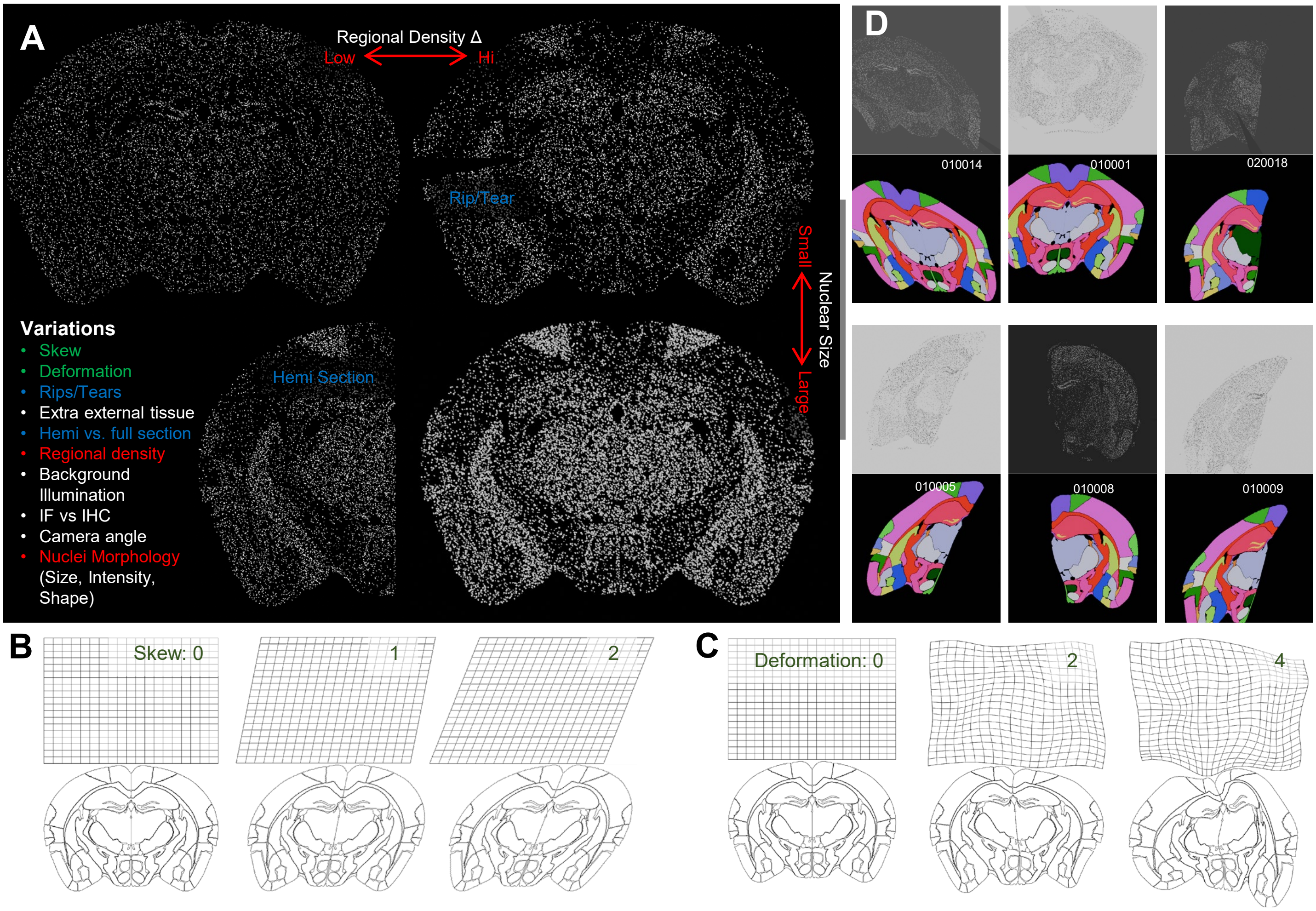


**Figure S11. Mouse Brain Macro simulation generation. A.** Overview of the different variations we can model using Blender Geometry Nodes. **B.** Skew moves the orientation of the plane of slice relative to the microscope, representing different slice angles, where 2 is a more extreme example. **C.** Deformation mimcs the natural variation observed in the shape of the slices between individuals. **D.** Examples of rendered macro images with different levels of variations with their respective labels. The white text is the image's unique ID.

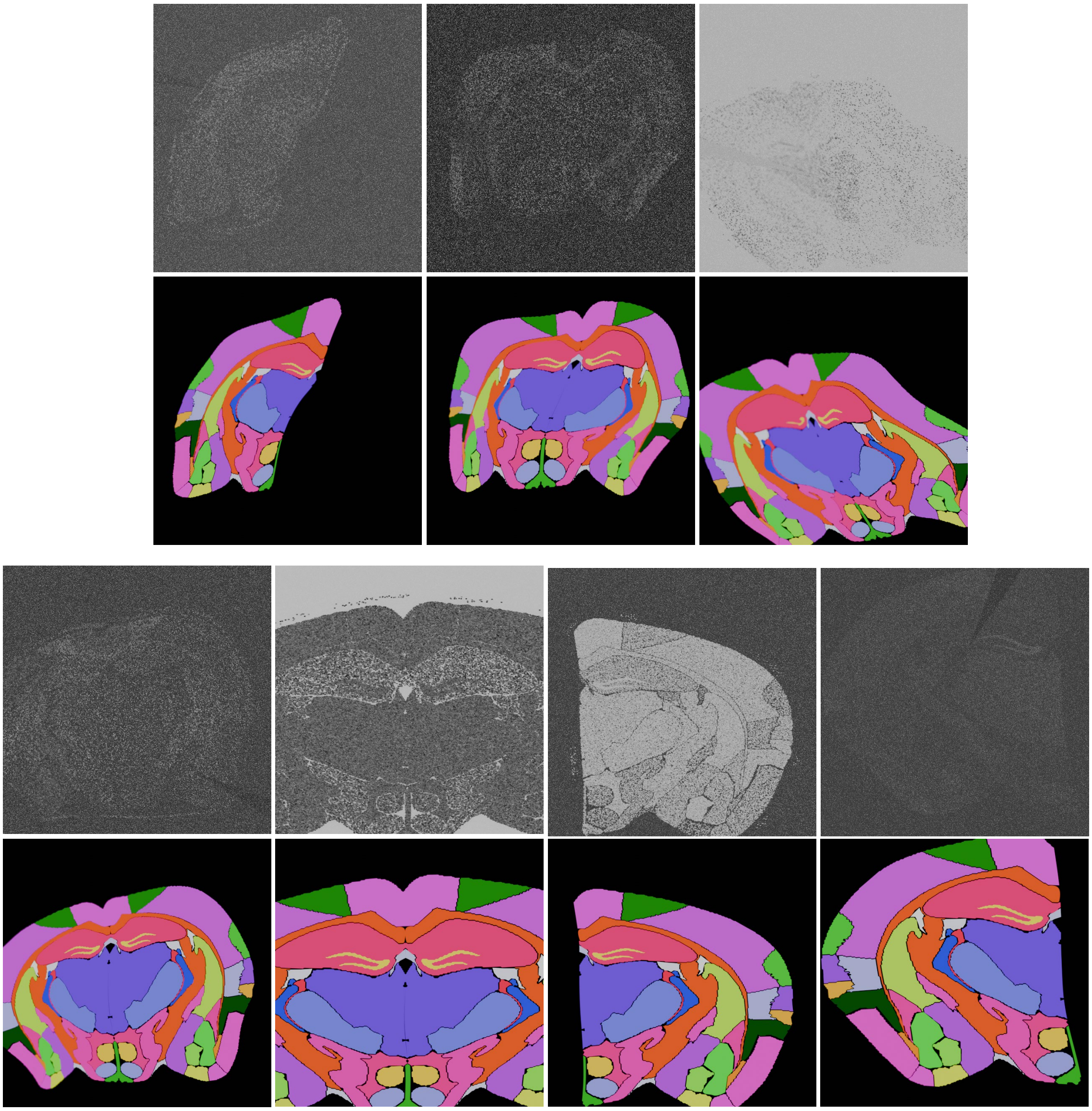

**Figure S12. Additional Training Examples.** Examples of cases of aberrations deliberately added to the brain macro image set to improve SiDoLa-NS-macro-mCB's generalizability. Demonstrated are examples of warp, skew, broken tissue, hemisection/full brain, light/dark background, and combinations of these features. The top row includes the model input images. The bottom row is the respective labels. Note that labels ignore the tissue aberrations such that the model learns to reconstruct the whole tissue.
